# The GPCR Smoothened on Cholinergic Interneurons Modulates Dopamine-associated Acetylcholine Dynamics and Affects Learning

**DOI:** 10.1101/2025.07.03.662982

**Authors:** Santiago Uribe-Cano, Andreas H. Kottmann

## Abstract

The striatum is a hub for associative learning where fluctuations in dopamine and acetylcholine dynamically regulate behavior. Acetylcholine is released by cholinergic interneurons, which integrate diverse inputs to contextualize dopamine signals and shape behavior. We previously observed that the GPCR Smoothened on cholinergic interneurons suppresses L-DOPA-induced dyskinesias, a motor side-effect resulting from medication elevated dopamine in the Parkinsonian brain. Here, we examine whether Smoothened signaling modulates acetylcholine dynamics, its coordination with dopamine, and motor learning in the healthy brain. We find that cholinergic neuron-specific Smoothened activity bidirectionally modulates acetylcholine inhibition following dopaminergic or cholinergic neuron activity. These effects alter the temporal organization of acetylcholine in the dorsolateral striatum and its coupling to dopamine. Behaviorally, Smoothened ablation from cholinergic neurons promotes motor learning and altered adjustment to changes in the effort or time to obtain reward. These findings identify Smoothened as a bidirectional modulator of striatal dopamine-acetylcholine coordination and striatal learning.

**Highlights:** Sonic Hedgehog–Smoothened signaling in striatal cholinergic interneurons bidirectionally regulates dopamine-associated cholinergic pauses without altering dopamine release.

Cholinergic Smoothened shapes the timing, duration, and coordination of endogenous dopamine–acetylcholine dynamics in the dorsolateral striatum.

Cholinergic Smoothened modulates striatal learning by accelerating motor learning while constraining effort management in an instrumental task.

**Graphical Abstract:** 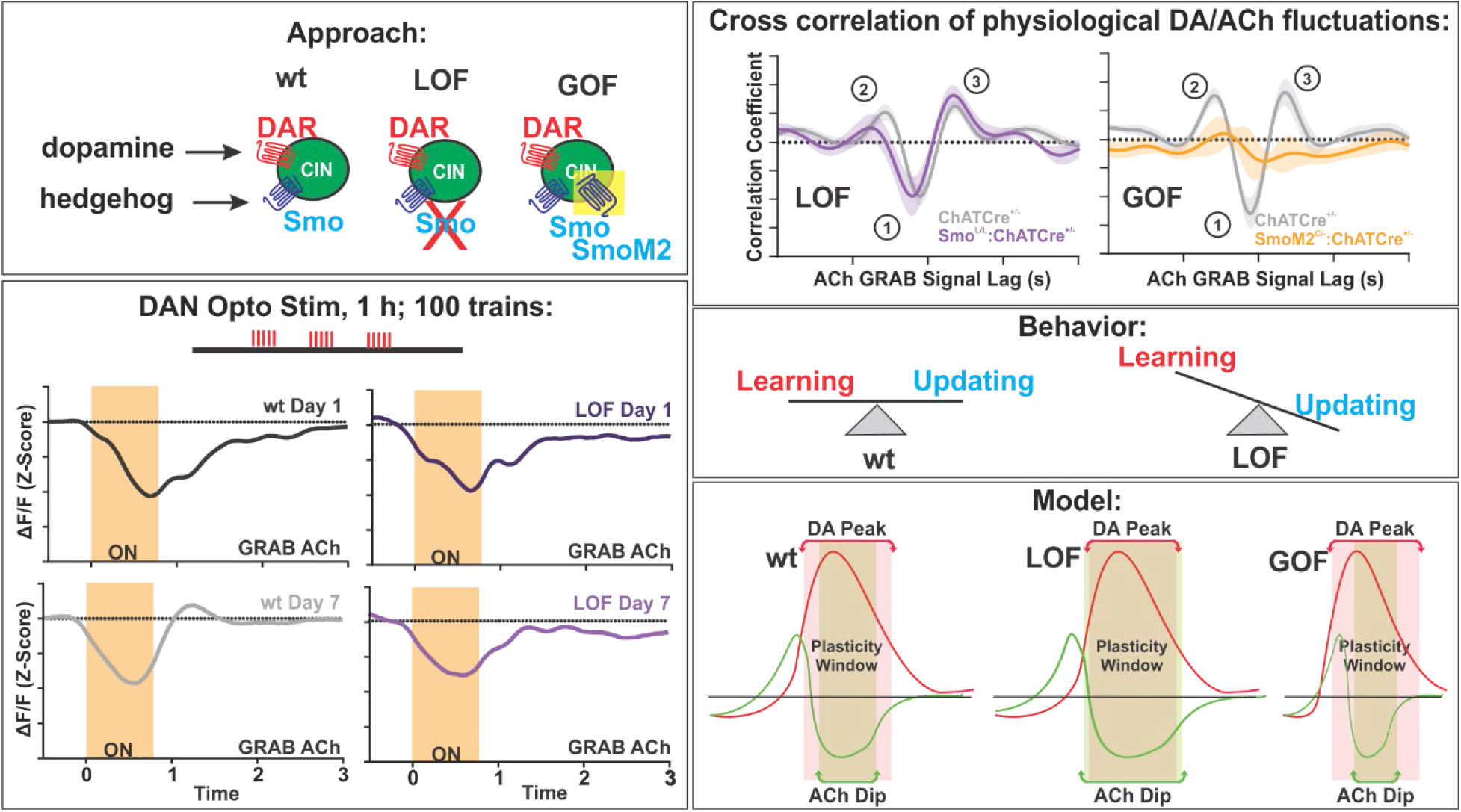

## Introduction

Cholinergic interneurons (CIN) of the striatum play a critical role in action selection, associative learning, and behavioral flexibility.^1–9^ While recognized for their tonic pacemaker-like activity in the 5–10 Hz range, CIN in the dorsolateral striatum (DLS) also exhibit multi-phasic responses consisting of sustained pauses often flanked by bursts of activity in response to salient stimuli.^7,10–13^ These “conditioned pauses” ^14,15^ are elicited by salient stimuli and develop during reinforcement learning in a manner anticorrelated with striatal dopamine (DA) release.^7,16^

The molecular mechanisms underlying the conditioned pause have received considerable attention, in part because the resulting reductions in acetylcholine (ACh) are thought to create a temporal window during which DA can modify corticostriatal synapses to drive learning.^7,12,17–19^ While glutamatergic input strength, intrinsic electrophysiological properties, GABAergic inhibition, and DA signaling have all been implicated in shaping CIN pauses (reviewed in ^20^, CIN receive input from over 40 neuromodulatory systems, including several peptidergic factors, many of which remain poorly characterized.^20,21^ This complex signaling landscape likely further tunes CIN output and downstream striatal function.^20–24^

Among these lesser-understood pathways, the secreted peptide Sonic Hedgehog (Shh) and its downstream effector, the GPCR Smoothened (Smo), appear to be critical modulators of CIN function. At long time-scales, Shh signaling on CIN appears critical for survival of CIN in mice ^25,26^, a finding supported by human data from familial and sporadic cases of Parkinson’s disease (PD).^27–31^ At more rapid time-scales relevant for real-time modulation of behavior, prior work shows that Shh signaling on CIN suppresses L-DOPA-induced dyskinesias (LID) ^32^, an aberrant form of motor learning associated with pathological CIN activity and caused by chronic levodopa treatment.^33^ These observations raise the question of whether and how Shh signaling contributes to CIN function in the healthy adult brain, particularly during periods of elevated DA, and whether it plays a role in coordinating DA-ACh interactions that underlie plasticity and motor learning.

Shh is expressed in the striatum by DAN and cortical layer 5 pyramidal tract (PTN) afferents, and can be released in an activity-dependent manner.^26,34–37^ Its receptor Patched (Ptch) is expressed in the striatum by CIN, Fast-Spiking Interneurons, and glia, ^26^ where it maintains cholesterol-dependent repression of Smo until it is bound by Shh (reviewed in ^38–40^). Both Ptch and Smo localize to the primary cilium and postsynaptic compartments ^38,41,42^, where Smo activation can trigger acute Gαi-coupled signaling and slower Gli-dependent transcriptional changes.^38,43,44^

Here, using cholinergic neuron-specific gain and loss of function manipulations, we reveal that Smo activity in the DLS bidirectionally modulates cholinergic inhibition, organizes DA-ACh dynamics, and impacts both motor and reinforcement learning.

## RESULTS

### Smo signaling on CIN modulates cholinergic inhibition following repeated dopaminergic stimulation

We previously showed that administration of Smo agonist (SAG) increases activity marker p-rpS6 in CIN and attenuates dyskinesias in a dose dependent manner.^32^ This suggested an acute, functionally opposing interaction between Smo and DA receptor signaling on CIN. Given that D2 receptor signaling prolongs the length of CIN pauses ^23,24,45,46^, we hypothesized that Smo signaling may counteract DA-mediated CIN inhibition. To test this, we manipulated Smo signaling and measured ACh levels following repeated optogenetic DA release in the DLS, a region critical for LID development ^47^ and known for robust DA-associated CIN pauses.^16^

Control or Smo^L/L^:ChATCre^+/-^ mice with cholinergic neuron-specific Smo ablation received AAV-Chrimson into the substantia nigra pars compacta (SNpc) and AAV expressing G protein-coupled receptor-based acetylcholine or dopamine sensors (GRAB ACh or GRAB DA) into the DLS, along with an optic fiber implant allowing simultaneous Chrimson stimulation and GRAB sensor photometry (Figures 1A and 1B). Despite notable spread of AAV-Chrimson expression at the level of the midbrain (Figure 1B; right), we found that expression within the striatum was highly localized to Tyrosine Hydroxylase (TH) positive fibers, indicative of selective local expression among DAN projections in the region (Figure 1B left; Figures 1A–1D). Specifically, the observed degree of correlation between TH and Chrimson expression in the striatum significantly exceeded that expected from 90% overlap between the two signals (Figure S1E; see figure legend for details), suggesting that together with the anatomical specificity of the fiber optic implant placement (Figure 1B, left), our approach stimulated primarily DAN terminals in the recording region.

**Figure 1.**
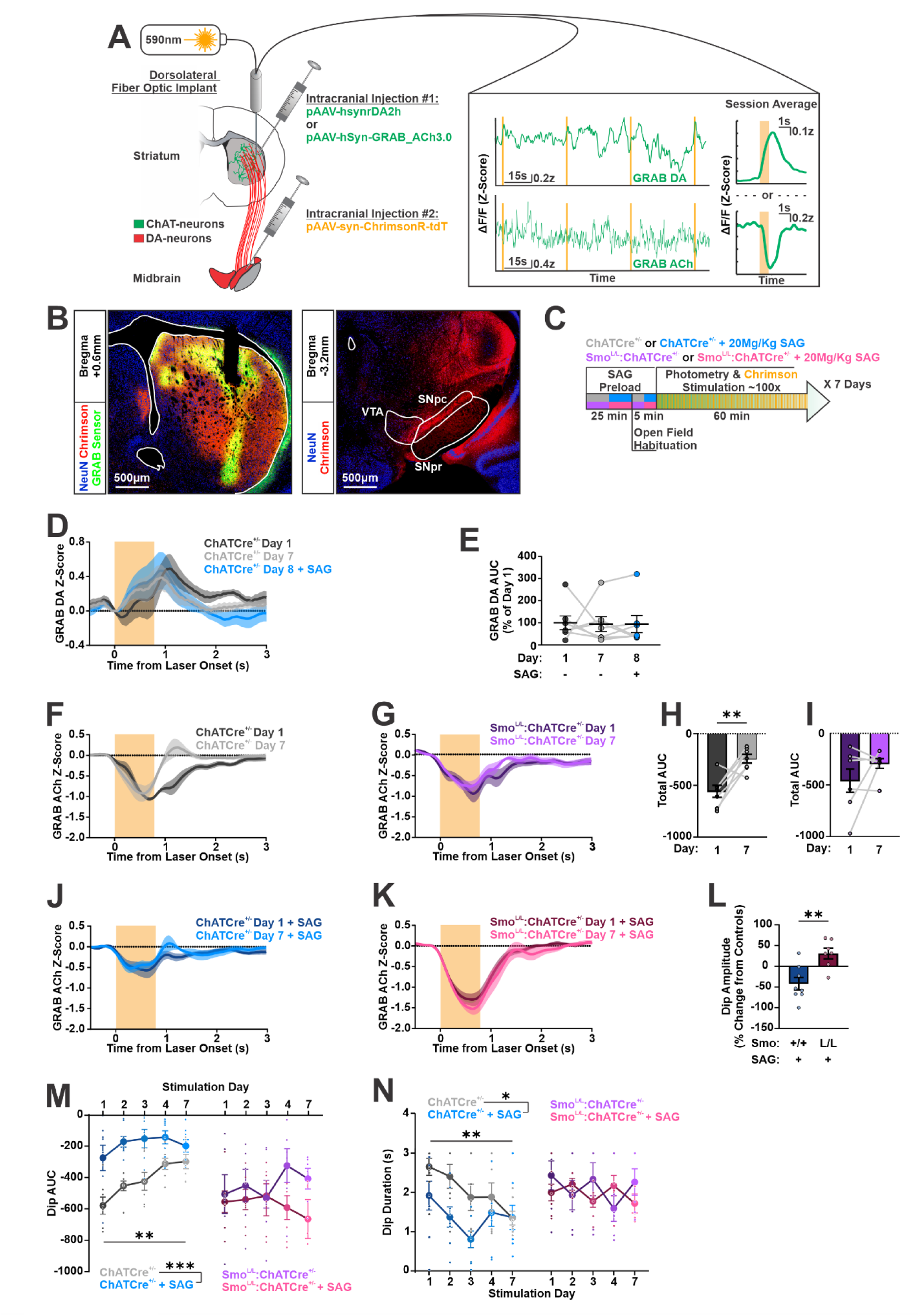
Smo on CIN modulates ACh inhibition following repeated DAN stimulation. **(A)** *Left:* Viral strategy for simultaneous dopamine neuron (DAN) axon terminal stimulation and G protein-coupled receptor-based sensor for dopamine (GRAB DA) or acetylcholine (GRAB ACh) recording in the dorsolateral striatum. *Right:* Representative traces across four laser pulse trains with session averages. **(B)** GRAB sensor and ChrimsonR-tdTomato expression in the dorsolateral striatum and substantia nigra pars compacta. Scale bar = 500µm. **(C)** Daily stimulation protocol in ChATCre^+/-^ or Smo^L/L^:ChATCre^+/-^ mice with or without Smoothened agonist (SAG). **(D)** Dopamine (DA) signals aligned to laser onset on days 1 and 7 in ChATCre^+/-^ mice (n = 7). A single SAG dose at 20Mg/Kg was administered on day 8 to assess effects on DA release. **(E)** Quantification of DA release area under the curve (AUC) from Panel (D) reported as percent of day 1 (n = 7; repeated-measures one-way ANOVA: F(1.378, 8.269) = 0.03, p > 0.05). For Supp Fig 1 **(F)** ACh signal on days 1 and 7 in ChATCre^+/-^ mice (n = 7). **(G)** Same as Panel (F), in Smo^L/L^:ChATCre^+/-^ mice (n = 6). **(H)** Comparison of GRAB ACh AUC on days 1 and 7 in ChATCre^+/-^ mice (n = 7 per day; paired two-tailed Student’s t test, **p < 0.01). **(I)** Comparison of GRAB ACh AUC on days 1 and 7 in Smo^L/L^:ChATCre^+/-^ mice (n = 6 per day; paired two-tailed Student’s t test, p > 0.05). **(J–K)** ACh signal in SAG-treated mice on days 1 and 7 (n = 7–8 per genotype). **(L)** Percent change in ACh dip amplitude in SAG-treated animals relative to untreated control average for each genotype (n = 6–7; paired two-tailed Student’s t test; p < 0.01). **(M)** ACh dip AUC across stimulation days with or without SAG treatment in ChATCre^+/-^ (n = 7–8 per day; two-way repeated-measures ANOVA: Day effect, F(2.902, 37.72) = 6.4, p < 0.01; Pharmacology effect, F(1, 13) = 23.47, p < 0.001; Day × Pharmacology interaction, F(4, 52) = 2.03, p > 0.05) and Smo^L/L^:ChATCre^+/-^ mice (n = 6–7 per day; two-way repeated-measures ANOVA: Day effect, F(2.615, 28.76) = 0.57, p > 0.05; Pharmacology effect, F(1, 11) = 1.41, p > 0.05; Day × Pharmacology interaction, F(4, 44) = 2.18, p > 0.05). **(N)** ACh dip duration across stimulation days with or without SAG treatment in ChATCre^+/-^ (n = 7–8 per day; two-way repeated-measures ANOVA: Day effect, F(3.491, 45.39) = 4.81, p < 0.01; Pharmacology effect, F(1, 13) = 5.52, p < 0.05; Day × Pharmacology interaction, F(4, 52) = 1.62, p > 0.05) and Smo^L/L^:ChATCre^+/-^ mice (n = 6–7 per day; two-way repeated-measures ANOVA: Day effect, F(2.785, 30.64) = 0.35, p > 0.05; Pharmacology effect, F(1, 11) = 0.57, p > 0.05; Day × Pharmacology interaction, F(4, 44) = 1.64, p > 0.05).

During an hour-long session, mice that had undergone surgery freely explored an open field while DAN terminals were repeatedly stimulated (Figure 1C). Based on prior evidence that Smo signaling in CIN modulates striatal plasticity across repeated DAN stimulation sessions ^32^, we repeated this procedure over seven consecutive days to assess how ACh release adapted over time.

In control animals expressing the GRAB DA sensor, optogenetic stimulation consistently evoked time-locked increases in DA that coincided with laser onset and remained stable in size across the seven sessions (Figure 1D and 1E). Similarly, Smo^L/L^:ChATCre^+/-^ mice expressing GRAB DA sensor also showed time-locked increases in DA release that showed no significant change from day 1 to day 7 (Figures S1F and S1G).

In a separate cohort of control mice expressing the GRAB ACh sensor, stimulation on day 1 elicited a robust ACh dip that began with laser onset and persisted for several seconds after laser offset (Figure 1F; black). Notably, this dip required Chrimson expression in the midbrain (Supplementary Figures 1H and 1I). After 7 consecutive daily sessions of stimulation, this ACh dip was significantly smaller compared to the first day and now included a rebound almost immediately following laser offset (Figure 1F; grey).

This shift from inhibition towards rebound was supported by a positive shift in the overall area under the curve (AUC) of the observed ACh response to stimulation (Figure 1H), a change reminiscent of CIN “conditioned pauses” previously described in the DLS and behaviorally linked to striatal learning^10,16,24,48^.

In contrast to controls, Smo^L/L^:ChATCre^+/-^ mice showed similar ACh dips on day 1 and day 7 with no evident rebound forming after the dip (Figure 1G). This lack of change was reflected by quantification of the overall AUC which showed no change between day 1 and day 7 (Figure 1I), revealing that Smo on CIN is required for the shift from inhibition to rebound observed among controls.

### Acute Smo activation on CIN modulates progressive changes in cholinergic inhibition

These initial experiments relied on germline mutations to manipulate Smo expression on CIN. While this approach is cholinergic neuron-specific, it leaves open the possibility that the observed effects result from secondary adaptations or developmental disturbances. To test whether acute Smo signaling modulates CIN inhibition, we repeated the stimulation protocol using control mice treated with Smoothened agonist (SAG) 30 minutes prior to each stimulation session (Figure 1C).

A challenge dose of SAG did not affect Chrimson-evoked DA release in controls (Figure 1D and 1E) or Smo^L/L^:ChATCre^+/-^ mice (Figure S1G). However, when compared to their own respective untreated counterparts on day 1, SAG treatment of ChATCre^+/-^ mice reduced the amplitude of ACh dips (Figures 1J and 1L) while SAG treatment of Smo^L/L^:ChATCre^+/-^ mice, with no Smo expression on CIN, increased the amplitude of ACh dips (Figures 1K and 1L). Further, repeated SAG treatment in controls over the course of 7 days of DAN stimulation appeared to qualitatively allow rebound formation while blunting the progressive reduction in dip size observed among untreated ChatCre^+/-^controls (Figure 1J). In contrast, and as was the case in untreated Smo^L/L^:ChATCre^+/-^ mice, repeated SAG treatment of mice lacking Smo expression on CIN produced comparable ACh dip sizes on days 1 and 7 of DAN stimulation with no evident rebound formation (Figure 1K).

To quantify these differences while also capturing potential progressive, Smo-dependent changes in specific aspects of the ACh profiles, we measured the total AUC, amplitude, and duration of ACh dips and subsequent rebounds, along with the time to the minimum of the dip or maximum of the rebound across all recording days (Figures 1M and 1N; Figure S2). We found that the overall size of ACh dips was reduced across days of stimulation in controls, suggesting that repeated DAN stimulation drives progressive reductions of ACh dip size (Figure 1M, grey). SAG treatment also further reduced dip size across all days (Figure 1M; grey vs blue), supporting the acute attenuation of CIN inhibition by SAG treatment. In contrast, rebound size showed no consistent changes following either repeated stimulation or SAG treatment (Figure S2A; grey vs blue), suggesting the overall shift towards reduced inhibition we observed in controls (Figure 1H) was driven by reductions in ACh dip size.

Consistent with the observed changes in AUC, we found that both stimulation day and SAG treatment reduced dip duration (Figure 1N) while SAG attenuated dip amplitude (Figure S2B; grey vs blue) and stimulation day attenuated time from laser onset to dip minimum (Figure S2C; grey vs blue). Notably, while the amplitude and duration of ACh rebounds were unaffected by SAG (Figures S2D and S2E; grey vs blue), the timing of the maximum rebound was decreased both across stimulation days and by SAG treatment (Figure S2F; grey vs blue), likely reflecting shorter ACh dips preceding rebound onset.

When we repeated these analyses with SAG-treated and untreated Smo^L/L^:ChATCre^+/-^ mice, we observed a lack of effect on all analyzed features (Figures 1M and 1N; Figure S2; red vs purple). Specifically, neither SAG treatment nor repeated days of DAN stimulation altered the size of these features in treated and untreated Smo^L/L^:ChATCre^+/-^ mice.

Together, these results indicate that concurrent Smo activity in CIN is required for the progressive reduction of CIN inhibition that occurs in response to repeated optogenetic DAN stimulation. In addition, the opposing effect on dip amplitude of SAG treatment in controls and Smo^L/L^:ChATCre^+/-^mice suggests that the absence of Smo signaling from CIN unmasks an effect of SAG on other striatal cell types, likely fast-spiking interneurons or astrocytes, which are known to engage in Smo signaling.^26^ The sum of those SAG effects on non-CIN cells might exert an inhibitory, secondary, effect on CIN that opposes the direct Smo-dependent modulation observed in controls.

### Smo manipulations impact ACh release associated with spontaneous DA release

Optogenetically evoked DAN activity can elicit non-physiological responses in target cells, raising concerns about how closely these manipulations reflect natural conditions.^49^ To address this, we developed an approach to detect endogenous “DA Events,” which presumably correspond to periods of DAN burst firing, as well as their coincident ACh signals in freely fluctuating GRAB DA and GRAB ACh traces from mice. To this end, we head-fixed Smo^L/L^:ChATCre^+/-^ mice and control Smo^L/+^:ChATCre^+/-^ littermates co-infused with a mix of AAVs expressing green-fluorescent GRAB ACh and red-fluorescent GRAB DA sensors in the DLS (Figures 2A and 2 B). After recovery and habituation, animals underwent a 30-minute recording session without any explicit behavioral task during which both signals were simultaneously recorded (Figure 2C).

**Figure 2.**
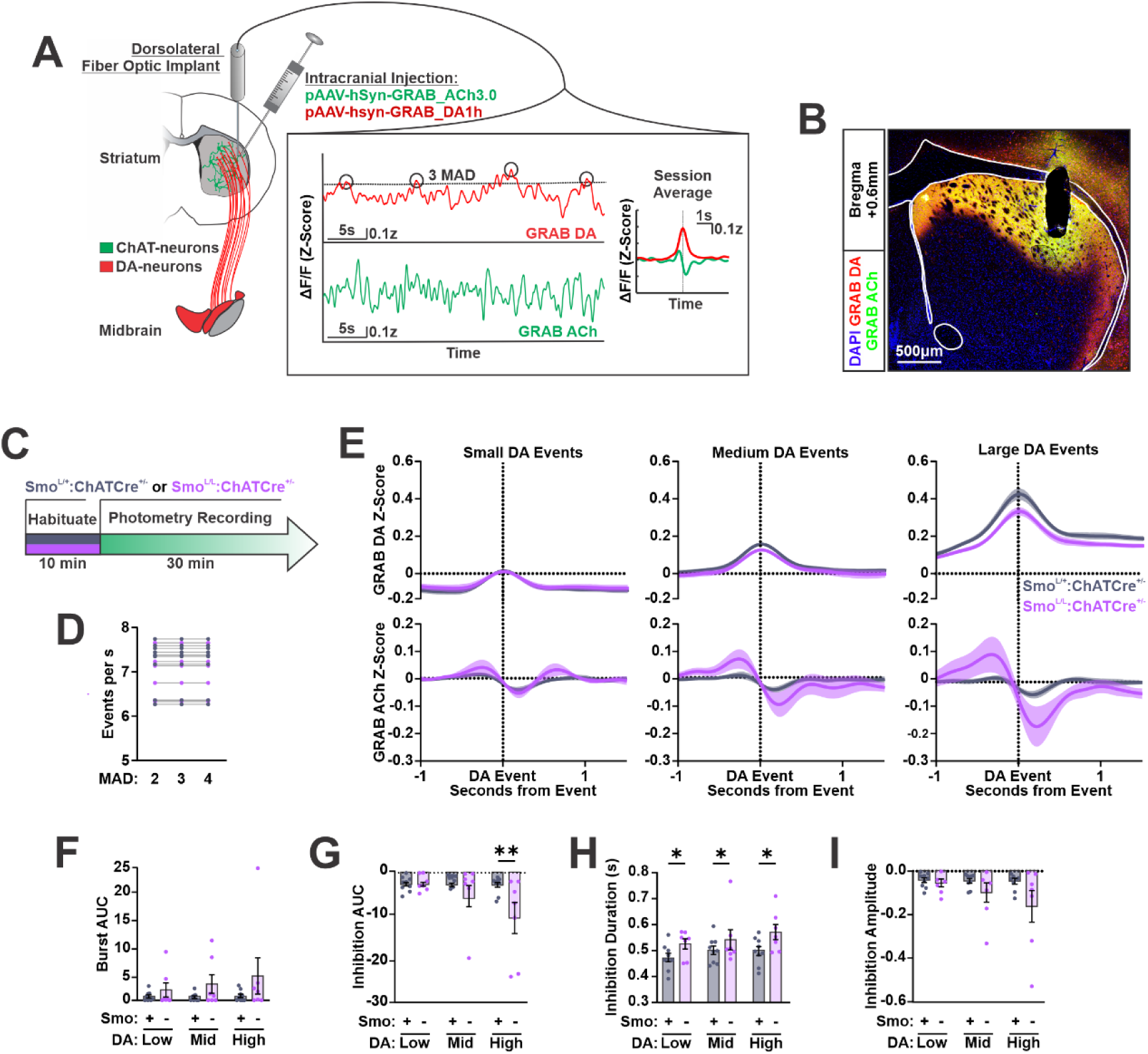
Smo on CIN modulates ACh inhibition coincident with endogenous DAN bursts. **(A)** *Left:* Viral strategy for simultaneous G protein-coupled receptor-based acetylcholine and dopamine sensor (GRAB ACh and GRAB DA) recordings in the dorsolateral striatum. *Right:* Representative traces from a single mouse, with session-averaged GRAB DA Event-aligned ACh profiles. **(B)** GRAB sensor expression in the striatum. Scale bar = 500µm. **(C)** Recording protocol for control and Smo^L/L^:ChATCre^+/-^ mice. **(D)** Frequency of detected DA Events with detection threshold set to 2, 3, or 4 MAD (n = 16 per MAD threshold). **(E)** Average Spontaneous DA Events stratified by amplitude (*top*) and coincident ACh signal (*bottom*) from control and Smo^L/L^:ChATCre^+/-^ mice (n = 7–9 per condition). **(F)** Area under the curve (AUC) for ACh bursts preceding DA events in Panel (D) (n = 7–9 per condition; two-way repeated measures ANOVA: Genotype effect, F(1, 14) = 2.07, p > 0.05; Coincident DA effect, F(1.143, 16.01) = 1.02, p > 0.05; Genotype × Coincident DA interaction, F(2, 28) = 0.84, p > 0.05). **(G)** AUC for ACh inhibition following DA events in Panel (D) (n = 7–9 per condition; two-way repeated measures ANOVA: Genotype effect, F(1, 14) = 3.22, p > 0.05; Coincident DA effect, F(1.555, 21.77) = 5.46, p < 0.05; Genotype × Coincident DA interaction, F(2, 28) = 5.80, p < 0.01; post hoc Šídák’s multiple comparisons test: **p < 0.01). **(H)** ACh inhibition duration in Panel (D) (n = 7–9 per condition; two-way repeated measures ANOVA: Genotype effect, F(1, 14) = 4.70, p < 0.05; Coincident DA effect, F(1.864, 26.09) = 2.02, p > 0.05; Genotype × Coincident DA interaction, F(2, 28) = 0.43, p > 0.05). No post hoc tests were performed due to the absence of a significant interaction effect; graph annotations denote significant main effects only. Inhibition duration was quantified on a trial-by-trial basis, capturing variability in event timing not reflected in group averages displayed in Panel (E). **(I)** ACh inhibition amplitude in Panel (D) (n = 7–9 per condition; two-way repeated measures ANOVA: Genotype effect, F(1, 14) = 2.42, p > 0.05; Coincident DA effect, F(1.370, 19.17) = 3.39, p > 0.05; Genotype × Coincident DA interaction, F(2, 28) = 2.94, p > 0.05).

To identify spontaneous increases in DAN activity we applied a 3-median absolute deviation (MAD) threshold to the GRAB DA signal and defined suprathreshold transients as “DA Events”. Although this threshold is arbitrary, it reliably captured transients consistent with experimenter judgment and has been both successfully used for the detection of calcium transients ^50^ and published in established fiber photometry analysis pipelines.^51^ Modifying this threshold to a level of 2 or 4 MAD did not alter the number of detected events for any animal (Figure 2D), suggesting this method robustly detects peaks in the photometry data without biases introduced by the specific threshold.

All DA Events extracted using this approach were aligned, along with their coincident ACh traces (see Methods), to generate session-averaged profiles for each signal. These averages revealed transient DA peaks of various magnitudes that were accompanied by biphasic ACh responses resembling burst-pause responses. Specifically, the DA Event-associated ACh profiles were characterized by increased ACh preceding the DA peak and decreased ACh following the DA peak (Figure 2A; Session Average). To assess how relatively different levels of DA might modulate the effect of Smo on ACh inhibition identified using this approach, we stratified DA Events into terciles representing the largest, middle, and smallest DA transients and then analyzed their associated ACh profiles (Figure 2E).

We found that coincident ACh bursts preceding DA Events did not differ significantly between Smo^L/L^:ChATCre^+/-^ mutants and controls regardless of DA event magnitude (Figure 2F). In contrast, the subsequent phase of ACh inhibition following DA Events of the top tercile was significantly larger among Smo^L/L^:ChATCre^+/-^ mice compared to controls (Figure 2G). This increase in overall inhibition was primarily driven by a genotype-dependent prolongation of inhibition duration, which was evident across all three terciles of DA Event magnitude (Figure 2H) but not by inhibition amplitude (Figure 2I).

These findings centered on endogenous increases in DA accord well with optogenetic results (Figure 1) by highlighting modulation of the duration of DA-associated ACh inhibition as a key feature of Smo’s action on CIN.

### Smo on CIN bidirectionally modulates cholinergic inhibition following ACh burst events

Inhibition of tonic CIN activity can occur due to cholinergic auto-inhibition, GABAergic input from other striatal neurons, or as a result of afterhyperpolarizations elicited by glutamatergic input from the cortex and thalamus. ^9,52–54^ These mechanisms however, and in particular glutamatergic input onto CIN, may not always be associated with coincident DA release events.^48^ Thus, restricting our analysis to ACh responses coincident with DA release could bias our findings toward a subset of all CIN inhibitory responses.

To address this, we modified our event detection pipeline to identify periods of elevated ACh that may reflect CIN burst activity, which are typically followed by afterhyperpolarizations but not necessarily associated with simultaneous DA release.^48^ Moreover, because Smo signaling can produce graded pathway activation depending on the level of receptor activity ^38^, we performed these experiments in both conditional loss- and gain-of-function Smo models anticipating opposing effects on CIN physiology. To study the impact of cholinergic-specific Smo loss-of-function, we continued using Smo^L/L^:ChATCre^+/-^and Smo^L/+^:ChATCre^+/-^ mice controls. For cholinergic-specific Smo gain-of-function, we utilized SmoM2^C/-^:ChATCre^+/-^ mice carrying a conditional allele of the constitutively active Smo variant, SmoM2, and ChATCre^+/-^ mice controls.

Applying the 3-MAD threshold to the GRAB ACh signal revealed spontaneous increases in cholinergic activity, which we extracted alongside coincident DA signals (Figure 3A). Despite expected variability across individual ACh-DA pairs, session-averaged traces revealed a consistent motif: ACh Events were flanked by transient dips in ACh, preceded by reduced DA, and followed by increased DA (Figure 3A; Session Average). This pattern reflects the well-documented inverse relationship between phasic ACh and DA in the striatum and fits the assumption that our detection method identifies, at least in part, CIN burst-firing events displaying subsequent afterhyperpolarizations.^48^

**Figure 3.**
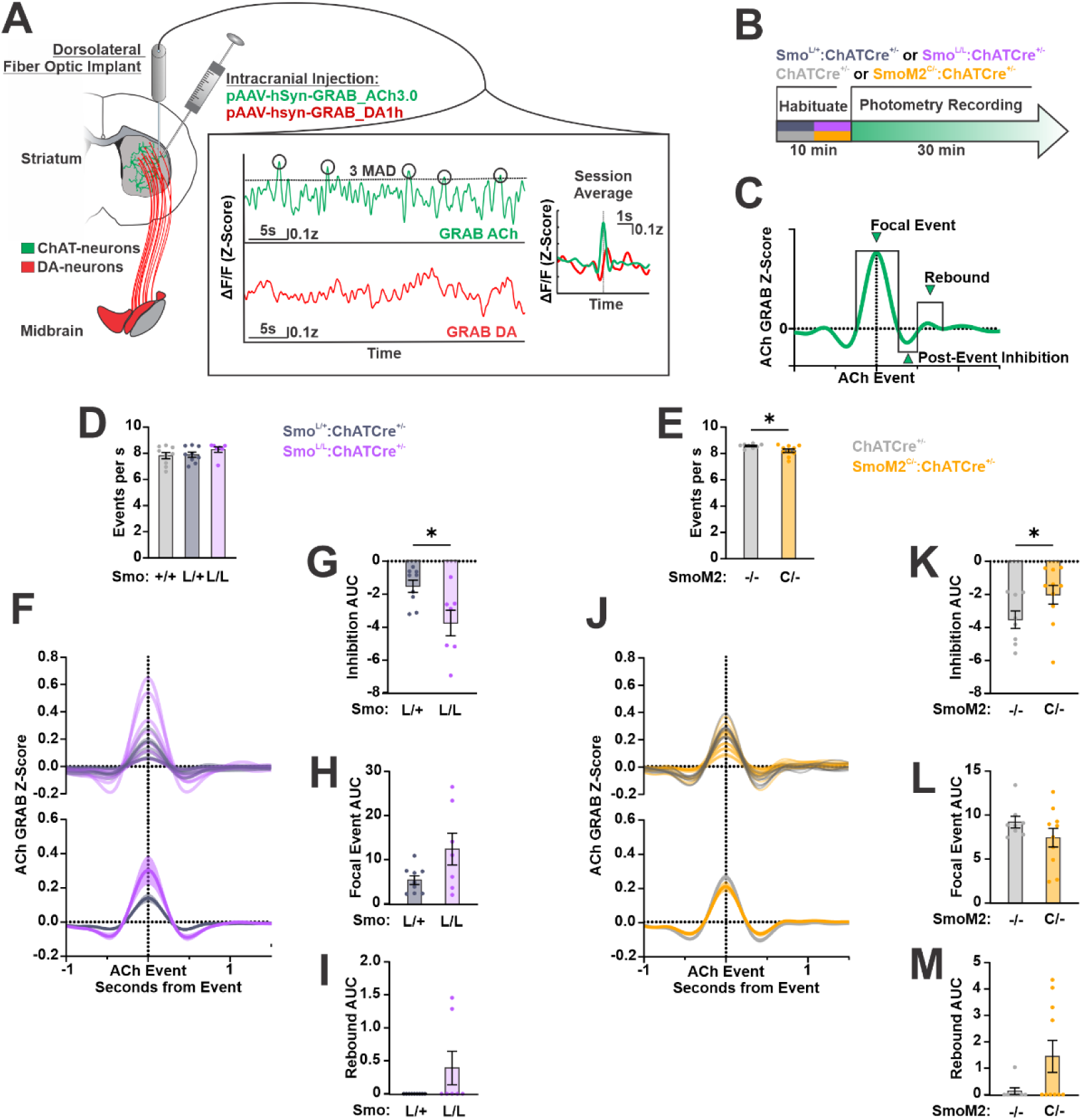
Smo on CIN bidirectionally modulates inhibition associated with spontaneous ACh Events. **(A)** *Left:* Viral strategy for simultaneous G protein-coupled receptor-based acetylcholine and dopamine sensor (GRAB ACh and GRAB DA) recordings in the dorsolateral striatum. *Right:* Representative traces from a single mouse, with session-averaged GRAB ACh Event-aligned profiles. **(B)** Recording protocol for control, Smo^L/L^:ChATCre^+/-^, and SmoM2^C/-^:ChATCre^+/-^ mice. **(C)** Schematic of quantified ACh Event features. **(D)** Frequency of detected ACh Events in Smo^L/L^:ChATCre^+/-^ mice versus controls (n = 7–9; Shapiro-Wilk test indicated lack of normality, p < 0.05; Kruskal-Wallis test, p > 0.05). **(E)** Frequency of detected ACh Events in SmoM2^C/-^:ChATCre^+/-^ mice versus controls (n = 8–10; F test indicated unequal variances, p < 0.05; Welch-corrected unpaired two-tailed t test, *p < 0.05). **(F)** Individual replicates and genotype-grouped acetylcholine (ACh) Events for controls and Smo^L/L^:ChATCre^+/-^ mice (n = 7–9 per genotype). **(G)** AUC for ACh Post-Event Inhibition from Smo^L/L^:ChATCre^+/-^ mice and controls in Panel (F) (n = 7–9 per genotype; unpaired two-tailed Student’s t test, *p < 0.05). **(H)** Area under the curve (AUC) for ACh Focal Events from Smo^L/L^:ChATCre^+/-^ mice and controls in Panel (F) (n = 7–9 per genotype; F test indicated unequal variances, p < 0.01; Welch-corrected unpaired two-tailed t test, p > 0.05). **(I)** AUC for ACh Rebound from Smo^L/L^:ChATCre^+/-^ mice and controls in Panel (F) (n = 7–9 per genotype; Shapiro-Wilk test indicated lack of normality, p < 0.001; F test indicated unequal variances, p < 0.0001; Mann-Whitney U test, p > 0.05). **(J)** Individual replicates and genotype-grouped ACh Events for controls and SmoM2^C/-^:ChATCre^+/-^ mice (n = 8–10 per genotype). **(K)** AUC for ACh Post-Event Inhibition from SmoM2^C/-^:ChATCre^+/-^ mice and controls in Panel (J) (n = 8–10 per genotype; Shapiro-Wilk test indicated lack of normality, p < 0.05; Mann-Whitney U test, *p < 0.05). **(L)** AUC for ACh Focal Events from SmoM2^C/-^:ChATCre^+/-^ mice and controls in Panel (J) (n = 8–10 per genotype; unpaired two-tailed Student’s t test, p > 0.05). **(M)** AUC for ACh Rebound from SmoM2^C/-^:ChATCre^+/-^ mice and controls in Panel (J) (n = 8–10 per genotype; Shapiro-Wilk test indicated lack of normality, p < 0.01; F test indicated unequal variances, p < 0.001; Mann-Whitney U test, p > 0.05).

To examine differences in ACh release between Smo mutants and their respective controls, we quantified the frequency of detected events as well as individual aspects of the average ACh Event profile. Specifically, we segmented the ACh profile into a Focal Event, Post-Event Inhibition, and Rebound phase (Figure 3C; see Methods). Loss-of-function Smo^L/L^:ChATCre^+/-^ mice showed no difference in event rate relative to heterozygous or wild-type controls (Figure 3D), whereas SmoM2^C/-^ :ChATCre^+/-^ gain-of-function mice exhibited a modest reduction in event frequency (Figure 3E).

When we performed a comparison of session-averaged ACh Event profiles between controls and loss-of-function Smo^L/L^:ChATCre^+/-^ mice (Figure 3F), we observed a significant increase in the AUC of the Post-Event Inhibition feature (Figure 3G), with no significant difference in the AUC of Focal Event or Rebound features (Figures 3H and 3I). Conversely, cholinergic-specific expression of SmoM2 (SmoM2^C/-^ :ChATCre^+/-^) produced the opposite effect (Figure 3J): a significant reduction in the AUC of average Post-Event Inhibition relative to controls (Figure 3K) with no significant changes in the Focal Event or Rebound (Figures 3L and 3M). Together, these results reveal that Smo bidirectionally modulates cholinergic pauses that follow spontaneous ACh burst events.

### The effect of Smo on CIN inhibition is strongest during periods of elevated DA levels

To better understand how coincident DA levels might influence the bidirectional effects of Smo on CIN inhibition, we ranked all detected ACh Events by their associated DA levels and compared averages derived from the top, middle, and bottom terciles of coincident DA (Figures 4A and 4H; see Figure 3S for illustration of the ranking and regrouping procedure). This process revealed that ACh Events could be observed across different levels of coincident DA (Figure 4A and 4H). Analysis of these stratified ACh Events revealed that the largest difference in Post-Event Inhibition AUC between control and Smo^L/L^:ChATCre^+/-^ mice occurred among ACh Events associated with the highest levels of coincident DA (Figure 4B). Consistent with our bulk analysis of unstratified ACh Events (Figure 3), no genotype differences were observed in the AUC of Focal Events or Rebounds across any of the DA terciles (Figures 4C and 4D).

**Figure 4.**
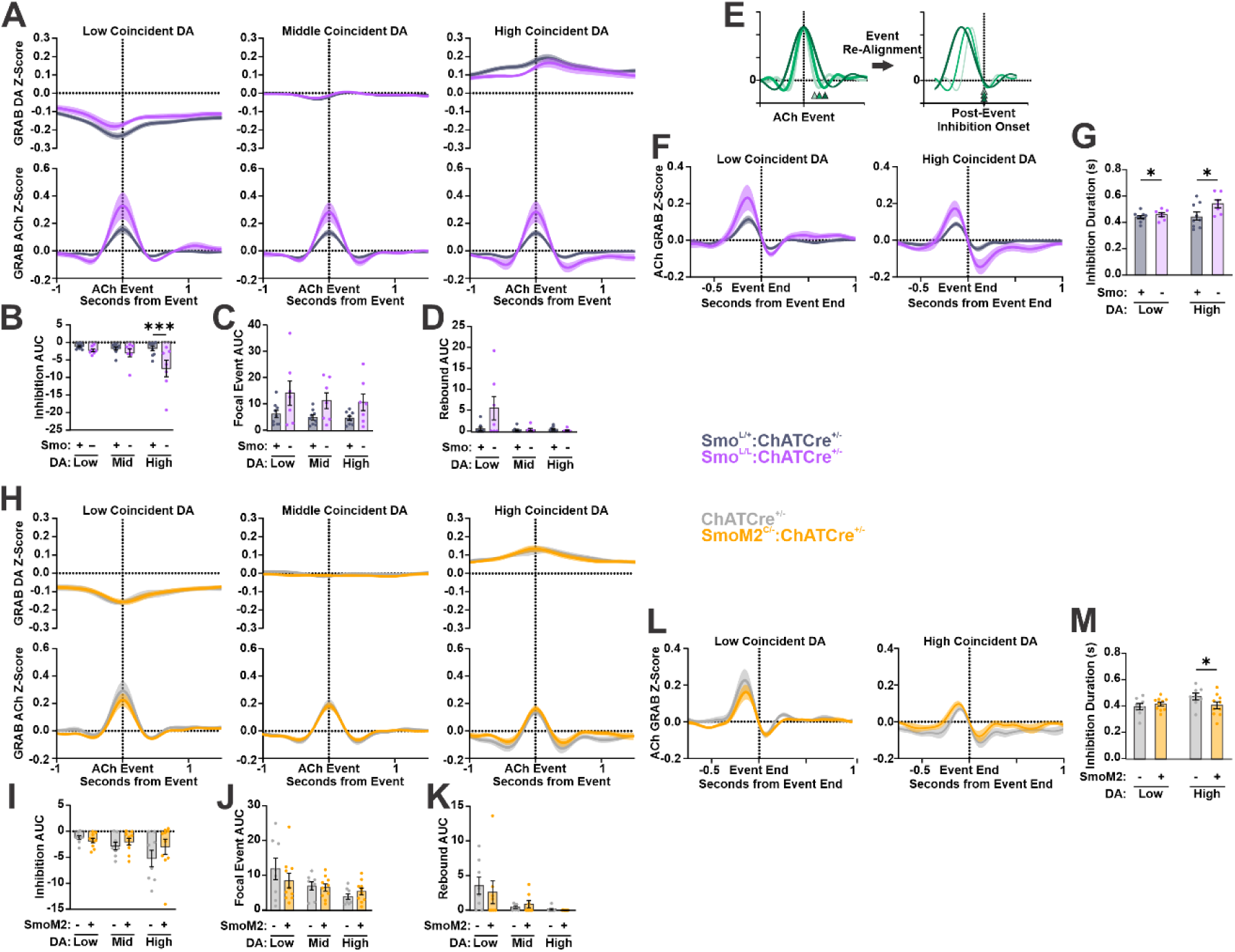
Bidirectional effects of Smo on CIN inhibition are most robust under high DA conditions. **(A)** Average ACh Events (*bottom*) grouped by coincident DA (*top*) levels from control and Smo^L/L^:ChATCre^+/-^ mice (n = 7–9 per condition). **(B)** Area under the curve (AUC) for ACh Post-Event Inhibition in Panel (A) (n = 7–9 per condition; two-way repeated measures ANOVA: Genotype effect, F(1, 14) = 6.48, p < 0.05; Coincident DA effect, F(1.397, 19.56) = 5.98, p < 0.05; Genotype × Coincident DA interaction, F(2, 28) = 4.63, p < 0.05; post hoc Šídák’s multiple comparisons test: ***p < 0.001). **(C)** AUC for ACh Focal Events in Panel (A) (n = 7–9 per condition; two-way repeated measures ANOVA: Genotype effect, F(1, 14) = 4.42, p > 0.05; Coincident DA effect, F(1.266, 17.72) = 3.40, p > 0.05; Genotype × Coincident DA interaction, F(2, 28) = 0.48, p > 0.05). **(D)** AUC for ACh Rebound in Panel (A) (n = 7–9 per condition; two-way repeated measures ANOVA: Genotype effect, F(1, 14) = 3.89, p > 0.05; Coincident DA effect, F(1.042, 14.59) = 4.49, p > 0.05; Genotype × Coincident DA interaction, F(2, 28) = 3.87, p < 0.05). **(E)** Schematic showing realignment of ACh Events to Post-Event Inhibition onset. **(F)** Control and Smo^L/L^:ChATCre^+/-^ ACh Events from Panel (A) realigned to Post-Event Inhibition onset (n = 7–9 per condition). **(G)** Duration of Post-Event Inhibition extracted from traces in Panel (F) (n = 7–9 per condition; two-way repeated-measures ANOVA: Genotype effect, F(1,14) = 6.01, p < 0.05; Coincident DA effect, F(1, 14) = 3.28, p > 0.05; Genotype × Coincident DA interaction, F(1,14) = 2.22, p > 0.05). No post hoc tests were performed due to the absence of a significant interaction effect; graph annotations denote significant main effects only.” **(H)** Average ACh Events (*bottom*) grouped by coincident DA (*top*) levels from control and SmoM2^C/-^:ChATCre^+/-^ mice (n = 8–10 per condition). **(I)** AUC for ACh Post-Event Inhibition in Panel (H) (n = 8–10 per condition; two-way repeated measures ANOVA: Genotype effect, F(1, 16) = 0.58, p > 0.05; Coincident DA effect, F(1.204, 19.27) = 6.1, p < 0.05; Genotype × Coincident DA interaction, F(2, 32) = 1.74, p > 0.05). **(J)** AUC for ACh Focal Events in Panel (H) (n = 8–10 per condition; two-way repeated measures ANOVA: Genotype effect, F(1, 16) = 0.20, p > 0.05; Coincident DA effect, F(1.051, 16.82) = 7.66, p < 0.05; Genotype × Coincident DA interaction, F(2, 32) = 1.44, p > 0.05). **(K)** AUC for ACh Rebound in Panel (H) (n = 8–10 per condition; two-way repeated measures ANOVA: Genotype effect, F(1, 16) = 0.08, p > 0.05; Coincident DA effect, F(1.092, 15.29) = 6.66, p < 0.05; Genotype × Coincident DA interaction, F(2, 28) = 0.33, p > 0.05). **(L)** Control and SmoM2^C/-^:ChATCre^+/-^ ACh Events from Panel (H) realigned to Post-Event Inhibition onset (n = 8–10 per condition). **(M)** Duration of Post-Event Inhibition extracted from traces in Panel (L) (n = 8–10 per condition; two-way repeated-measures ANOVA: Genotype effect, F(1,16) = 0.90, p > 0.05; Coincident DA effect, F(1, 16) = 2.50, p > 0.05; Genotype × Coincident DA interaction, F(1,16) = 4.51, p < 0.05; post hoc Šídák’s multiple comparisons test: *p < 0.05).

As in our optogenetic experiments, Smo ablation affected ACh inhibition through changes in both amplitude (Supplementary Figure 4A) and duration (Figure 4G). When ACh Events from the top and bottom terciles were realigned to the onset of Post-Event Inhibition, a clear visual difference in both these measures emerged between the genotypes under high- but not low-coincident DA conditions (Figures 4E and 4F). However, when quantified, these differences remained a genotype main effect rather than an interaction (Figure 4G; Figure S4A) and there was no effect on the timing of the dip minimum after the focal event (Figure S4B), suggesting that unlike the overall AUC measure of Post-Event Inhibition (Figure 4B), these individual measures did not rely on coincident DA levels.

The use of heterozygous Smo^L/+^:ChATCre^+/-^ controls in these experiments was chosen to ensure stringent genetic matching and control for potential effects of the floxed Smo allele. To further validate our findings, we expanded the analysis to include ChATCre^+/-^ controls carrying two intact Smo alleles (Figure S5). Incorporating these additional controls again underscored a consistent genotype difference in the duration of ACh Post-Event Inhibition across levels of coincident DA, supporting that Smo manipulations alter the duration of ACh inhibition (Figure S5J).

When data from SmoM2^C/-^:ChATCre^+/-^ gain-of-function mice and controls was stratified by coincident DA level (Figure 4H), the main genotype effect observed in the Post-Event Inhibition AUC of unstratified ACh Events (Figure 3K) became less distinct, likely reflecting increased variance across DA terciles (Figure 4I). Focal Event AUC and Rebound AUC, as in Smo^L/L^:ChATCre^+/-^ mice, were not effected in SmoM2^C/-^:ChATCre^+/-^ (Figures 4J and K). Further, inhibition amplitude and timing of dip maximum from focal event was not effected (Figures S4C and S4D). However, analysis of Post-Event Inhibition duration again revealed the influence of Smo manipulations (Figure 4M). Remarkably, SmoM2-mediated gain-of-function in CIN opposed the Smo loss-of-function effect and instead decreased the duration of ACh inhibition for events associated with elevated DA levels (Figures 4L and 4M).

Together, these findings indicate that Smo ablation exaggerates CIN pauses, especially their duration, whereas constitutive SmoM2 activation shortens them. In addition, these effects do not appear to be exclusively linked to coincident DA levels.

### Multivariate analysis of changes in ACh signaling due to Smo on CIN

To fully assess the effect of Smo manipulations on ACh Event features, we performed principal component (PC) analysis on 51 parameters across all ACh Events from all mice, capturing information about feature size, rate of change, timing, and relationship to coincident DA (Supplementary Figure S6A and S6B).

In Smo^L/L^:ChATCre^+/-^ animals and controls, the largest source of variance (PC1; 19.73% variance) did not fully separate the two genotypes in PC space, as the distributions overlapped (Figure S6C). However, ACh Events from Smo^L/L^:ChATCre^+/-^ animals were shifted towards more negative PC1 values, resulting in a statistically significant difference in average PC1 value between genotypes (Figure S6C). A complementary result was observed in SmoM2^C/-^:ChATCre^+/-^ animals, where PC1 (17.26% variance) again did not fully differentiate the genotypes but showed a significant shift in average PC1 value due to reduced negative spread among SmoM2^C/-^:ChATCre^+/-^ samples (Figure S6D). These data suggest that Smo manipulations do not broadly reshape all ACh Events but rather affect a subset mainly concentrated in the negative PC1 range. This aligns with our observation that bidirectional Smo effects are most evident under certain conditions, namely, elevated DA.

To identify the features driving PC1 separation, we examined the highest PC1 loading factors (|loading| > 0.5) and isolated parameters common to both comparisons (Figure S6E). In both Smo^L/L^:ChATCre^+/-^ and SmoM2^C/-^:ChATCre^+/-^ datasets, variability in similar features, particularly the rate of ACh signal change and relative sizes of Post-Event Inhibition and Rebound components, contributed to the observed differences. Notably, the relative amplitude of the Post-Event Inhibition relative to the Focal Event (Parameter #23) showed the strongest association with PC1 (loading = −0.887) in both analyses, indicating that greater Post-Event Inhibition correlates with more negative PC1 values. This finding supports the observation that ACh Post-Event Inhibition, a feature that again maps onto CIN inhibition, is a key mechanism by which Smo signaling impinges on CIN.

### Smo contributes to the temporal organization of ACh-DA dynamics

Despite the anticorrelated and phase-shifted relationship of DA and ACh across behavioral states (Figure 2A and Figure 3A), DAN and CIN respond to largely distinct sets of inputs.^48^ Growing evidence, however, also indicates that mechanisms for localized crosstalk between DA and ACh exist within the striatum, implying their coordinated release is subject to precise regulation.^23,24,55^ We therefore examined whether Smo in CIN impinges on the temporal coordination of DA and ACh by computing cross-correlations between these signals in Smo loss- and gain-of-function mice relative to controls (Figures 5A and 5E).

**Figure 5.**
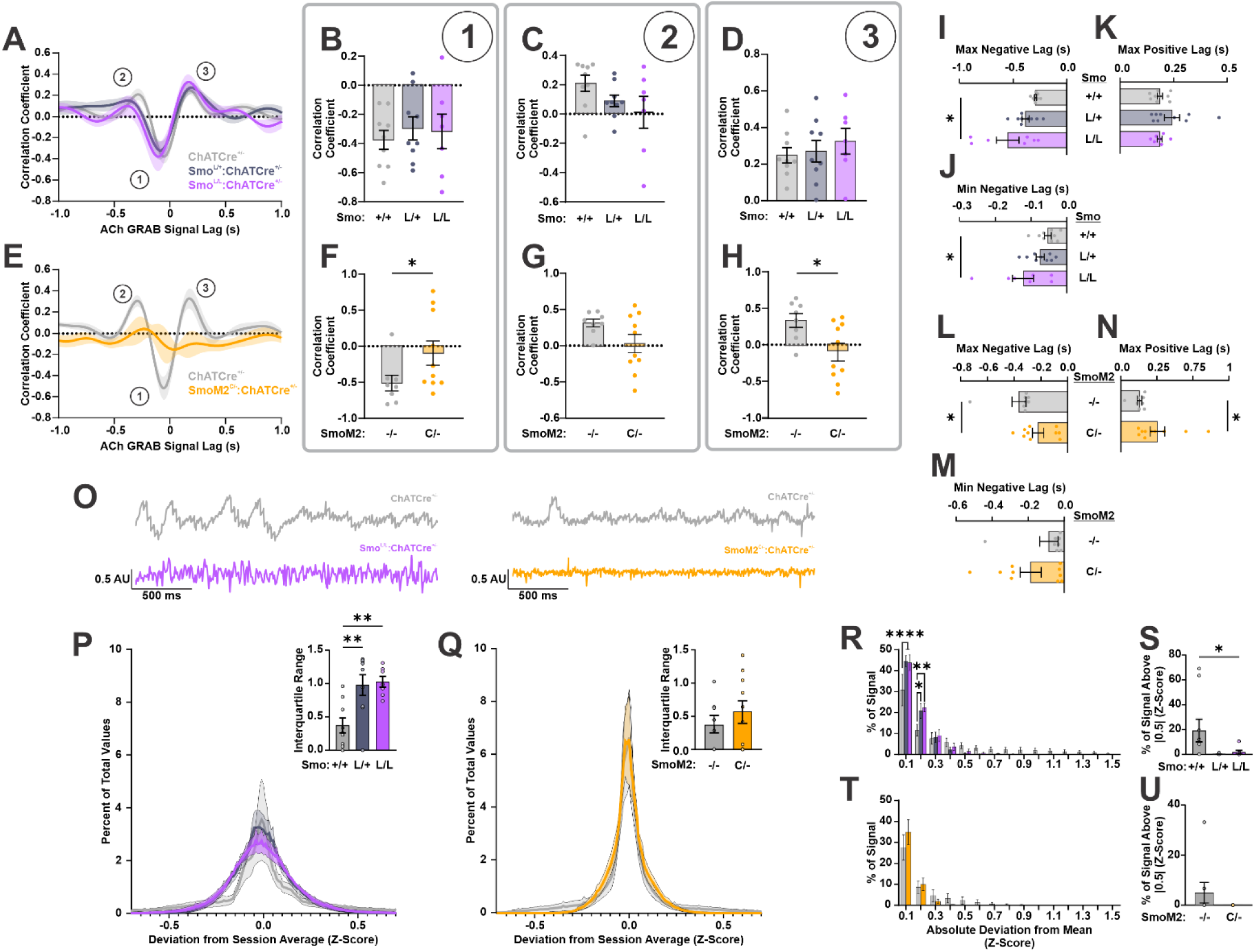
CIN Smo is required for normal ACh dynamics and DA coordination. **(A)** Average cross-correlation between acetylcholine (ACh) and dopamine (DA) signals aligned to ACh Events for Smo^L/L^:ChATCre^+/-^ mice and controls. Numbered features correspond to Panels (B–D) (n = 7–9 mice per genotype). **(B)** Quantification of correlation coefficient at feature 1 for controls and Smo^L/L^:ChATCre^+/-^ mice (n = 7–9 per genotype; one-way ANOVA: main effect, F(2, 22) = 0.24, p > 0.05). **(C)** Quantification of correlation coefficient at feature 2 for controls and Smo^L/L^:ChATCre^+/-^ mice (n = 7–9 per genotype; one-way ANOVA: main effect, F(2, 22) = 2.13, p > 0.05). **(D)** Quantification of correlation coefficient at feature 3 for controls and Smo^L/L^:ChATCre^+/-^ mice (n = 7–9 per genotype; one-way ANOVA: main effect, F(2, 22) = 0.45, p > 0.05). **(E)** Average cross-correlation between ACh and DA signals aligned to ACh Events for SmoM2^C/-^:ChATCre^+/-^ mice and controls. Numbered features correspond to Panels (F-H) (n = 8–10 mice per genotype). **(F)** Quantification of correlation coefficient at feature 1 for controls and SmoM2^C/-^:ChATCre^+/-^ mice (n = 8–10 per genotype; Shapiro-Wilk test indicated lack of normality, p < 0.05; Mann-Whitney U test, *p < 0.05). **(G)** Quantification of correlation coefficient at feature 2 for controls and SmoM2^C/-^:ChATCre^+/-^ mice (n = 8–10 per genotype; F test indicated unequal variances, p < 0.05, Welch-corrected unpaired two-tailed t test, p > 0.05). **(H)** Quantification of correlation coefficient at feature 3 for controls and SmoM2^C/-^:ChATCre^+/-^ mice (n = 8–10 per genotype; unpaired two-tailed Student’s t test, *p < 0.05). **(I)** Average lag for maximum correlation coefficient with a negative lag for controls and Smo^L/L^:ChATCre^+/-^ mice (n = 7–9 per genotype; Brown-Forsythe test indicated unequal variances, p < 0.01; Welch’s ANOVA test: main effect, F(2, 9.743) = 6.181, *p < 0.05). **(J)** Average lag for minimum correlation coefficient with a negative lag for controls and Smo^L/L^:ChATCre^+/-^ mice (n = 7–9 per genotype; one-way ANOVA on timing: main effect, F(2, 22) = 4.12, p < 0.05; post hoc Tukey’s multiple comparisons test: *p < 0.05). **(K)** Average lag for maximum correlation coefficient with a positive lag for controls and Smo^L/L^:ChATCre^+/-^ mice (n = 7–9 per genotype; Shapiro-Wilk test indicated lack of normality, p < 0.05; Kruskal-Wallis test, p > 0.05). **(L)** Average lag for maximum correlation coefficient with a negative lag for controls and SmoM2^C/-^:ChATCre^+/-^ mice (n = 8–10 per genotype; unpaired two-tailed Student’s t test, *p < 0.05). **(M)** Average lag for minimum correlation coefficient with a negative lag for controls and SmoM2^C/-^:ChATCre^+/-^ mice (n = 8–10 per genotype; unpaired two-tailed Student’s t test on timing, p > 0.05). **(N)** Average lag for maximum correlation coefficient with a positive lag for controls and SmoM2^C/-^:ChATCre^+/-^ mice (n = 8–10 per genotype; Shapiro-Wilk test indicated lack of normality, p < 0.01; F test indicated unequal variances, p < 0.01; Mann-Whitney U test, *p < 0.05). **(O)** Representative G protein-coupled receptor-based ACh sensor (GRAB ACh) traces. **(P)** Distribution of ACh values relative to the session average for Smo^L/L^:ChATCre^+/-^ and control mice. *Inset:* interquartile ranges (n = 7–9 per genotype; one-way ANOVA on interquartile range: main effect, F(2, 22) = 8.47, p < 0.01; post hoc Tukey’s multiple comparisons test: **p < 0.01). **(Q)** Distribution of ACh values relative to the session average for SmoM2^C/-^:ChATCre^+/-^ and control mice. *Inset:* interquartile ranges (n = 8–10 per genotype; unpaired two-tailed Student’s t test on interquartile range, p > 0.05). **(R)** Percent of ACh values binned by absolute deviation (0.1 z-score bins) for Smo^L/L^:ChATCre^+/-^ and control mice (n = 7–9 per genotype; two-way repeated-measures ANOVA: Genotype effect, F(2, 22) = 0.13, p > 0.05; Deviation effect, F(1.777, 39.10) = 95.2, p < 0.0001; Genotype × Deviation interaction, F(28, 308) = 2.45, p < 0.0001; post hoc Šídák’s multiple comparisons test: ****p < 0.0001, **p < 0.01, *p < 0.05). **(S)** Percent of ACh values with absolute deviation greater than 0.5 for Smo^L/L^:ChATCre^+/-^ and control mice (n = 7–9 per genotype; Shapiro-Wilk test indicated lack of normality, p < 0.0001; Brown-Forsythe test indicated unequal variances, p < 0.01; Kruskal-Wallis test, *p < 0.05). **(T)** Percent of ACh values binned by absolute deviation (0.1 z-score bins) for SmoM2^C/-^:ChATCre^+/-^ and control mice (n = 8–10 per genotype; two-way repeated-measures ANOVA: Genotype effect, F(1, 16) = 0.02, p > 0.05; Deviation effect, F(1.583, 25.33) = 39.65, p < 0.0001; Genotype × Deviation interaction, F(14, 224) = 0.86, p > 0.05). **(U)** Percent of ACh values with absolute deviation greater than 0.5 for SmoM2^C/-^:ChATCre^+/-^ and control mice (n = 8–10 per genotype; Shapiro-Wilk test indicated lack of normality, p < 0.0001; F test indicated unequal variances, p < 0.0001; Mann-Whitney U test, p > 0.05).

By focusing on ACh Events and their cross-correlation with coincident DA transients, this analysis revealed three prominent average-correlation peaks in controls (Figures 5A and 5E; grey): a negative one at −50ms (peak 1; indicating the extent to which DA dips predict ACh Events), and positive ones at −200ms (peak 2; indicating the extent to which DA release predicts ACh inhibition recovery) and +200ms (peak 3; indicating the extent by which ACh Events predict subsequent DA release). Comparison of Smo^L/L^:ChATCre^+/-^ loss-of-function mice to ChATCre^+/^ or Smo^L/+^:ChATCre^+/-^ controls revealed no difference in correlation magnitude at any of the three peaks (Figures 5B–5D), suggesting that the strength of DA–ACh coupling was not affected by Smo ablation from CIN.

In contrast, comparing gain-of-function SmoM2^C/-^:ChATCre^+/-^ mice to controls revealed that the strength of average correlations at control-defined peaks eroded in SmoM2^C/-^:ChATCre^+/-^ animals (Figure 5E) and was significantly reduced at peak 1 and peak 3 (Figures 5F–5H).

We next examined whether Smo manipulations altered the timing variability of these cross-correlation peaks. This analysis was motivated by the observation that peak-correlation timing showed substantial animal-to-animal dispersion, particularly within the mutant groups (Figures S7A–S7E). To formally assess differences in within-group variability, we applied Brown–Forsythe tests (a median-based test for equality of variances) to the observed timing of peak correlations from animals of each genotype. This analysis revealed significant differences in timing dispersion across genotypes, supporting the observation that Smo manipulations impacted within-group variability in the timing of peaks with negative lag (peak 1 and 2; Figures S7F and S7G) relative to controls. While still significant, this effect was less evident for the timing of peak-correlation with a positive lag (peak 3; Figure S7H). Thus, in order to better preserve timing structure at the level of individual ACh Event–DA pairs and avoid the temporal smoothing that might occur when cross-correlation waveforms are derived at the session level, we computed cross-correlations for each ACh Event–DA pair in a recording session and extracted the lag values corresponding to peaks 1–3. These lag values were then averaged across events within a session, yielding an animal-level mean peak timing measure.

This analysis revealed that the increased within-group-variability of peak-correlation timing was accompanied by shifts in average peak-correlation timing among Smo mutants. On average, both the minimum and the maximum correlations with negative lag (corresponding to peaks 1 and 2, respectively) occurred nearly twice as late in Smo^L/L^:ChATCre^+/-^ mice relative to ChATCre^+/-^ controls (Figures 5I and 5J), suggesting a delay in how DA fluctuations are mirrored by changes in ACh. This effect was gene dosage dependent, with heterozygous controls revealing an intermediate delay. In contrast, positive-lag maximum correlations (corresponding to peak 3) did not differ in timing between genotypes (Figure 5K). Thus, the loss of Smo from CIN selectively increases the lag of ACh responses to DA, without affecting the temporal relationship by which ACh predicts subsequent DA release.

In SmoM2^C/-^:ChATCre^+/-^ gain-of-function mice, the negative lag minimum did not shift significantly compared to controls (Figure 5M). However, the negative-lag maximum (corresponding to peak 2, Figure 5L) appeared sooner and the positive-lag maximum (corresponding to peak 3; Figure 5N) appeared later compared to controls. These findings indicate that SmoM2 expression on CIN shortens the delay by which CIN recovery follows DA release and increases the delay between CIN activity and subsequent DA release. Thus, constitutively active SmoM2 does not abolish DA–ACh coupling per se. Instead, like Smo^L/L^:ChATCre^+/-^ manipulations, it increases temporal variability. However, unlike the loss-of-function scenario, it shifts the direction of ACh–DA timing relationships in a manner that is partially opposite to the shift observed in Smo^L/L^:ChATCre^+/-^ mice.

### Smo modulates the dynamic range of striatal ACh

CIN exhibit intrinsic rhythmicity that sustains tonic ACh levels and allows fluctuations above and below baseline. Changes in CIN activity could therefore impact tonic ACh levels and change the dynamic range of potential ACh fluctuations. Given that Smo manipulations altered the duration of ACh inhibition across our analyses and experimental paradigms, we asked whether Smo also affects the overall dynamic range of the ACh signal beyond discrete ACh Events. To test this, we calculated the session-average of each 30-minute GRAB ACh recording and measured how much each sample deviated from that mean (Figures 5P and 5Q). We then plotted the percentage of samples across deviation magnitudes and compared distributions between mutants and controls (Figures 5R–5U).

In both Smo^L/L^:ChATCre^+/-^ and Smo^L/+^:ChATCre^+/-^ animals, we observed broader distributions around the mean ACh signal relative to ChATCre^+/-^ controls, reflected by a significantly increased interquartile range (Figure 5P). This broadening came at the expense of fewer high-deviation values at the tails, supported by binned quantification of absolute deviations (Figure 5R) and a reduced proportion of values > 0.5 z-scores from the mean (Figure 5S). SmoM2^C/-^:ChATCre^+/-^ mice showed a similar, though nonsignificant, trend (Figures 5Q, 5T and 5U), suggesting a potential ceiling on this effect in the gain-of-function scenario.

Together, these results suggest that endogenous fluctuations in Smo signaling within CIN are critical for maintaining the dynamic range of ACh activity, whereas static gain- or loss-of-function manipulations may constrain this range.

### Smo on CIN bidirectionally modulates motor learning

Given coordinated striatal DA and ACh transients are thought to create temporally defined windows for plasticity, our findings suggest that Smo manipulations may influence striatal learning. To test this, we assessed motor learning using the accelerating rotarod, a task previously shown to rely on striatal plasticity.^56^

Mice were trained over 8 consecutive days with 10 trials per day. Each trial began with an acceleration phase (0–40 RPM over 300 seconds), followed by a holding phase where top speed was maintained for 400 seconds. To isolate the learning component, we focused on the acceleration phase and defined “criterion” as the first training day on which mice reached top speed on > 50% of trials (Figure 6A). Learning rate was quantified as the slope of a linear regression fitted to each animal’s latency to fall on all trials up to and including the criterion day (Figure 6B). Performance during the holding phase was analyzed separately to assess potential effects on motor coordination or physical ability that were independent of learning.

**Figure 6.**
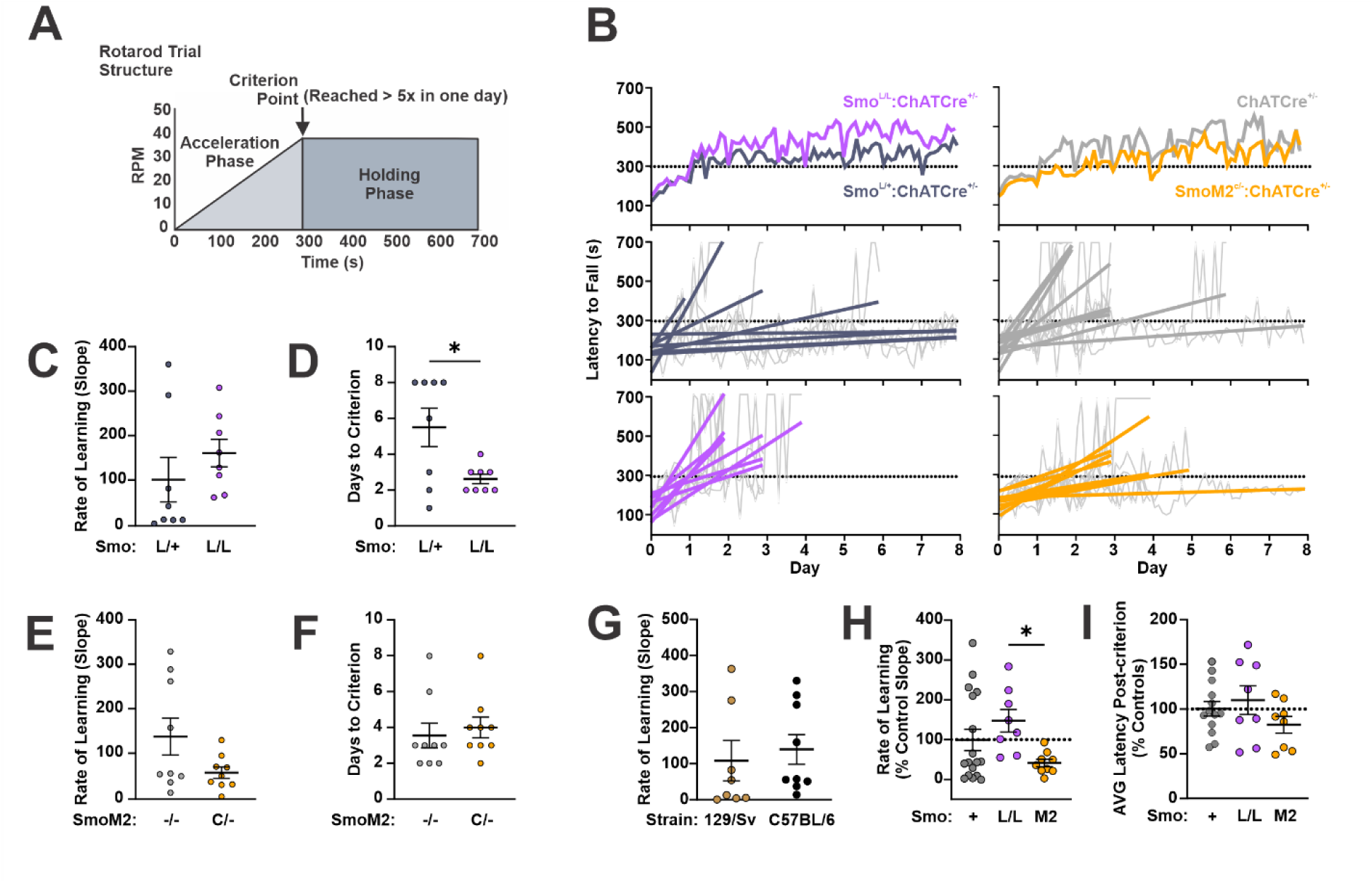
Smo bidirectionally impacts motor learning. **(A)** Structure of individual rotarod trials, performed 10 times per day. **(B)** Average latency to fall (*top*) and best-fit performance curves to criterion (*middle* and *bottom*) for Smo^L/L^:ChATCre^+/-^ (*left, bottom*) and SmoM2^C/-^:ChATCre^+/-^ (*right, bottom*) versus respective controls (middle). **(C)** Average slope of individual best-fit performance curves for Smo^L/L^:ChATCre^+/-^ and control mice (n = 8 per genotype; Shapiro-Wilk test indicated lack of normality, p < 0.01; Mann-Whitney U test, p > 0.05). **(D)** Average number of days to reach criterion for Smo^L/L^:ChATCre^+/-^ and control mice (n = 8 per genotype; Brown-Forsythe test indicated unequal variances, p < 0.01; Welch-corrected unpaired two-tailed t test, *p < 0.05). **(E)** Average slope of individual best-fit performance curves for SmoM2^C/-^:ChATCre^+/-^ and control mice (n = 9–10 per genotype; F test indicated unequal variances, p < 0.01; Welch-corrected unpaired two-tailed t test, p > 0.05). **(F)** Average number of days to reach criterion for SmoM2^C/-^:ChATCre^+/-^ and control mice (n = 9–10 per genotype; Shapiro-Wilk test indicated lack of normality, p < 0.01; Mann-Whitney U test, p > 0.05). **(G)** Slope comparison between control mouse strains (n = 8–9 per strain; Shapiro-Wilk test indicated lack of normality, p < 0.01; Mann-Whitney U test, p > 0.05). **(H)** Best-fit slopes normalized to background strain (n = 8–17 per genotype; Shapiro-Wilk test indicated lack of normality, p < 0.01; Kruskal-Wallis test, *p < 0.05; post hoc Dunn’s multiple comparisons test: *p < 0.05). **(I)** Average latency to fall post-criterion, normalized to background strain (n = 8–17 per genotype; one-way ANOVA: main effect, F(2, 26) = 1.36, p > 0.05).

Comparison of controls and loss-of-function Smo^L/L^:ChATCre^+/-^ animals revealed an increased learning rate in mutants (Figures 6B and 6C) confirmed by a significantly reduced number of days to reach criterion (Figure 6D). In contrast, gain-of-function SmoM2^C/-^:ChATCre^+/-^ animals exhibited a trend toward impaired learning, with a reduced learning rate that approached significance (Figure 6E; p = 0.09).

Since controls from the two background strains used to generate Smo^L/L^:ChATCre^+/-^ and SmoM2^C/-^:ChATCre^+/-^ animals showed no differences in baseline learning rate on the rotarod (Figure 6G), we pooled all controls and compared across genotypes to assess whether gain- and loss-of-function Smo manipulations exert opposing effects on motor learning. Indeed, when mutant performance was normalized to controls and directly compared, a bidirectional effect of Smo signaling on CIN during motor learning was observed (Figure 6H). Importantly, no differences were observed between genotypes in post-criterion performance (Figure 6I), suggesting differences in learning rate were not confounded by motor ability.

### Smo on CIN modulates striatal reinforcement learning

Instrumental reinforcement learning under variable interval (VI) schedules, during which rewards become available only after a variable delay, has been shown to depend on the DLS.^57^ To test whether Smo manipulation in CIN affects this form of learning, we trained Smo^L/L^:ChATCre^+/-^ and control animals on an escalating VI schedule (Figure 7A; see Methods) and assessed multiple aspects of instrumental behavior. Following three days of continuous reinforcement (CRF), animals underwent VI15 training for three days, followed by a reward devaluation probe to test behavioral flexibility. Training concluded with eight additional days of VI30 training.

**Figure 7.**
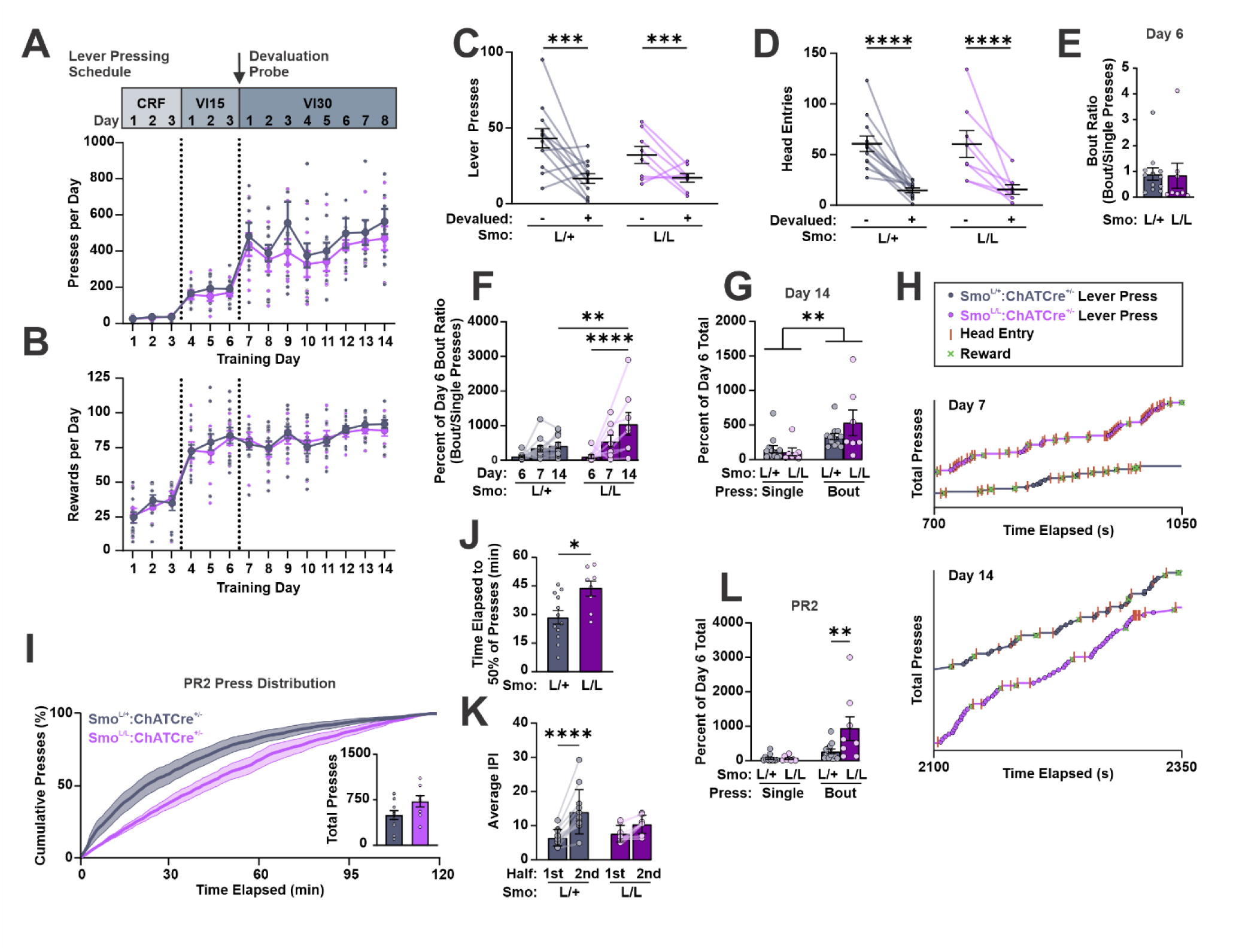
Smo ablation from CIN promotes pressing bouts and persistent instrumental responding. **(A)** Lever presses across initial days of continuous reinforcement (CRF) and subsequent days of escalating variable interval (VI) reinforcement (n = 8–12 per genotype; two-way repeated-measures ANOVA: Training day effect, F(4.47, 80.42) = 26.33, p < 0.0001; Genotype effect, F(1, 18) = 1.33, p > 0.05; Training day × Genotype interaction, F(13, 234) = 0.36, p > 0.05). **(B)** Rewards earned across VI training days (n = 8–12 per genotype; two-way repeated-measures ANOVA: Training day effect, F(6.765, 121.8) = 44.65, p < 0.0001; Genotype effect, F(1, 18) = 0.16, p > 0.05; Training day × Genotype interaction, F(13, 234) = 0.30, p > 0.05). **(C)** Lever presses with or without prior reward devaluation (n = 8–12 per condition; two-way repeated-measures ANOVA: Devaluation effect, F(1, 18) = 16.72, p < 0.001; Genotype effect, F(1, 18) = 1.06, p > 0.05; Devaluation × Genotype interaction, F(1, 18) = 1.26, p > 0.05). **(D)** Magazine entries with or without prior reward devaluation (n = 8–12 per condition; two-way repeated-measures ANOVA: Devaluation effect, F(1, 18) = 43.31, p < 0.0001; Genotype effect, F(1, 18) = 0.00, p > 0.05; Devaluation × Genotype interaction, F(1, 18) = 0.01, p > 0.05). **(E)** Ratio of press bouts to single presses performed on the final day of VI15 training (day 6). Higher values indicate a greater preference for bout pressing (n = 8–12 per genotype; Shapiro–Wilk test indicated non-normal distribution, p < 0.0001; Mann–Whitney U test, p > 0.05). **(F)** Change in bout ratio across days of VI30 training. Values are expressed as the percent of each animal’s ratio on day 6 (the final day of VI15 training). Higher values indicate a greater preference for bout pressing (n = 8–12 per condition; two-way repeated-measures ANOVA: Training day effect, F(1.339, 24.10) = 13.36, p < 0.001; Genotype effect, F(1, 18) = 4.36, p = 0.05; Training day × Genotype interaction, F(2, 36) = 3.29, p < 0.05; post hoc Šídák’s multiple comparisons test: ****p < 0.0001, **p < 0.01). **(G)** Number of single presses and press bouts performed on day 14, expressed as a percent of the respective values for each animal on day 6 (n = 8–12 per condition; two-way repeated-measures ANOVA: Press type effect, F(1, 17) = 12.54, p < 0.01; Genotype effect, F(1, 17) = 0.88, p > 0.05; Press type × Genotype interaction, F(1, 17) = 1.95, p > 0.05). **(H)** Example behavior traces showing lever presses, magazine entries, and rewards delivered across a span of 250–350 seconds on the first and last days of VI30 training. **(I)** Cumulative lever press distribution across the progressive ratio 2 (PR2) session. *Inset:* Total lever presses during the 2-hour session (n = 8–12 per genotype; unpaired two-tailed Student’s t test on total presses, p > 0.05). **(J)** Average latency to complete 50% of total lever presses (n = 8–12 per genotype; unpaired two-tailed Student’s t test, *p < 0.05). **(K)** Average inter-press interval (IPI) for each genotype, calculated separately for presses occurring in the first and second halves of the PR2 session (n = 8–12 per condition; two-way repeated-measures ANOVA: Session half effect, F(1, 18) = 33.89, p < 0.0001; Genotype effect, F(1, 18) = 0.57, p > 0.05; Session half × Genotype interaction, F(1, 18) = 7.36, p < 0.05; post hoc Šídák’s multiple comparisons test: ****p < 0.0001). **(L)** Number of single presses and press bouts performed during PR2 session, expressed as a percent of the respective values for each animal on day 6 (n = 8–12 per condition; two-way repeated-measures ANOVA: Press type effect, F(1, 18) = 4.59, p < 0.05; Genotype effect, F(1, 18) = 14.14, p < 0.01; Press type × Genotype interaction, F(1, 18) = 5.58, p < 0.05; post hoc Šídák’s multiple comparisons test: **p < 0.01).

Across the first 6 days of training, Smo^L/L^:ChATCre^+/-^ loss-of-function mutants and controls exhibited similar increases in lever pressing and earned comparable numbers of rewards as the reinforcement schedule escalated (Figures 7A and 7B). After the third day of VI15 training, animals were given one hour of free access to either the training reward (+devalued condition) or standard chow (-devalued condition), followed by a 15-minute extinction probe to assess devaluation sensitivity. As expected for goal directed learning associated with relatively limited training, animals significantly reduced lever pressing in the +devalued condition compared to the -devalued condition (Figure 7C). Both genotypes also showed reduced head entries into the reward magazine in the +devalued condition, supporting the efficacy of our devaluation protocol and suggesting a similar capacity between genotypes to modulate reward-directed behavior at this point in training (Figure 7D).

We next examined whether, despite similar total press counts and rewards earned, the two genotypes had developed distinct lever-pressing strategies at this stage of training. Mice tended to exhibit two approaches: either performing isolated lever presses followed by immediate head entries to check for rewards or executing grouped bouts of presses before checking the reward port. To quantify these behaviors, we calculated the ratio of press bouts to single presses exhibited by mice at this time point (day 6) and found no significant genotype differences, indicating that both groups had developed comparable lever-pressing behavior by this point (Figure 7E). This similarity allowed us to use each animal’s day 6 performance as a baseline going forward to assess how individual pressing strategies evolved with extended training.

An additional eight days of training were carried out under an escalated VI30 schedule. Although both genotypes continued to produce similar total press counts and earn comparable rewards (Figures 7A and 7B), Smo^L/L^:ChATCre^+/-^ mice displayed a significant increase in average bout ratio by the end of VI30 training, indicating a growing preference for press bouts over single presses (Figures 7F and 7H). To determine whether this shift reflected a reduction in single presses or an increase in bout pressing, we normalized the number of each press type on the final training day to that animal’s respective count on day 6. This analysis showed that the elevated bout ratio among mutants was primarily driven by an increase in press bouts from day 6 to day 14. Control animals also exhibited a modest increase in total press bouts, but this was offset by a similar rise in single presses that despite not reaching significance was sufficient to prevent a net change in bout ratio across training (Figures 7F and 7G).

CIN have been implicated in behavioral flexibility.^1,58^ The preference for bout pressing observed among Smo^L/L^:ChATCre^+/-^ mice could therefore reflect a deficit in action switching, such that mutants remain more persistently engaged at the lever and less frequently incorporate other behaviors such as checking the reward magazine or exploring the chamber.^1,58^ To test this, we evaluated mice that had undergone the same VI training up to day 12 in a Progressive Ratio (PR) task, where the number of presses required for each successive reward doubled, allowing direct assessment of task engagement. Although total press counts did not differ between groups, control mice tended to front-load their responses, performing most lever presses early in the session, whereas mutants displayed a more uniform response distribution across time (Figure 7I). This difference was confirmed by quantifying the average latency to reach 50% of total presses, which was shorter in controls (Figure 7J). Moreover, only control mice showed a significant difference in average inter-press interval between the first and second half of the session (Figure 7K), indicating that they reduced responding as the cost of rewards increased. In contrast, Smo^L/L^:ChATCre^+/-^ mice maintained a consistent response rate throughout, suggesting reduced sensitivity to evolving task contingencies and sustained engagement as session demands increased.

To determine whether the bout-preferring strategy observed during VI30 training persisted in the PR2 task, we compared the number of single presses and press bouts produced by each genotype across the entire session, normalized to the respective values on day 6 of training. Consistent with their VI30 performance, Smo^L/L^:ChATCre^+/-^ mice again exhibited a stronger preference for press bouts relative to controls (Figure 7L).

Together, these results suggest that the changes in ACh inhibition observed in Smo^L/L^:ChATCre^+/-^mice are associated with a behavioral phenotype characterized by accelerated motor learning, increased behavioral stereotypy, and reduced action switching. Future experiments incorporating direct measures of ACh dynamics during reversal or extinction learning will be critical for testing this proposed link between Smo-dependent modulation of ACh signaling and behavioral flexibility.

## Discussion

The processes that result in long-term corticostriatal plasticity in medium spiny projection neurons (MSNs) and enable learning are tightly governed by dynamic temporal and spatial patterns of extracellular DA and ACh.^59^ Recent studies have shown that similarly organized fluctuations of DA and ACh also occur in mice at rest, suggesting that the coordination of these neuromodulators during learning is enabled by an underlying circuit architecture that supports complex rhythms of DA and ACh.^48^ Here, focusing on CIN, we identify the atypical GPCR Smo, an effector of the Shh pathway, as a bidirectional modulator of the coordination between DA and ACh in the DLS of mice that are not engaged in an explicit behavioral task. Our results, based on conditional gain and loss function manipulations, suggest that Smo is a critical component of those basic circuit mechanism that enable coordinated DA and ACh rhythms.

Specifically, we found that enhanced Smo activity reduces the magnitude and duration of ACh inhibition during periods of elevated DA, whereas Smo loss prolongs inhibition and prevents the progressive shortening of the CIN pause that is otherwise observed in response to repeatedly evoked DA release. Smo manipulations also bidirectionally shifted the timing of ACh–DA correlations and constrained the dynamic range of ACh. These impacts of Smo on the organization of cholinergic activity patterns impinged on learning since we found that Smo ablation accelerated motor learning and promoted operant task engagement at the expense of dynamically updating action-outcome contingency values.

### Smo activity on CIN modulates the duration of ACh pauses in the DLS

ACh generally serves as an inhibitory signal in the striatum by suppressing corticostriatal transmission via presynaptic M2 receptors and by reducing MSN activity through GABAergic interneurons ^7,19,54,60,61^. Pauses in CIN activity, combined with the rapid ACh-clearing action of abundant Acetylcholinesterase (AChE) in the striatum, create transient ACh-free periods during which learning related changes in synaptic strengths can manifest.^7,19,62–64^ Specifically, for long-term corticostriatal plasticity on MSNs to occur, phasic release of DA and the pause in ACh must synchronize with glutamate-driven MSN depolarizations.^65^ Thus, under this model, processes that prolong the CIN pause increase the likelihood that synaptic inputs onto MSNs will be altered, whereas shortening the pause reduces it.

One factor known to influence the duration of the CIN pause in a bimodal manner is D2 receptor (D2R) activation. While D2R ablation from CIN shortens the pause, D2R overexpression on CIN extends it.^23,24^ Here, we find that Smo signaling on CIN also affects the cholinergic pause, but in a manner opposite to D2R: Elevated Smo signaling shortens the pause, whereas Smo ablation increases it. These results suggest that Smo might play a role in safeguarding against run away DA driven plasticity by supporting and possibly fine tuning the ability of CIN to gate and contextualize DA driven reinforcement signals.

### Behavioral correlates of altered Smo signaling on CIN

Our behavioral results begin to implicate specific behavioral changes as consequences of altered Smo signaling on CIN. In particular, we observed that loss of Smo from cholinergic neurons accelerated learning in a motor task and promoted a preference for repeated lever-pressing bouts in a progressive ratio task, suggesting reduced sensitivity to changing reward-outcome contingencies.

It is important to note that the gain- and loss-of-function models used in these behavioral experiments manipulated Smo across all cholinergic neurons. Thus, the observed behavioral effects could arise from changes to CIN activity within striatal sub-compartments beyond the DLS. For example, prior studies have demonstrated that manipulations to CIN activity within the dorsomedial striatum (DMS) affect reversal and extinction learning, ^9,66^ two cognitive processes that could account for some of the behavioral features observed here. However, unlike our findings, those studies ruled out changes in the cholinergic pause as a mechanism underlying their effects. These differences may reflect distinct yet parallel processing roles of the DLS and DMS, a likely scenario supported by the different patterns of coordinated DA and ACh observed in each compartment.^16,45,67^

One consideration, given the model described above in which Smo signaling modifies the width of the window of temporal integration at corticostriatal synapses, is that Smo-dependent modulation of cholinergic pauses may influence sequential learning in particular by altering the effective lifetime of eligibility traces at synapses representing sequences of actions marked for plasticity.^68^ Consistent with this idea, animals with cholinergic-specific Smo ablation favored pressing bouts over single presses in an operant task, supporting the interpretation that prolonged cholinergic pauses might allow stronger reinforcement of more temporally distant sequential actions. Alternatively, an expanded plasticity window may enhance behavioral acquisition without changes in temporal integration by reducing opposition to DA-mediated plasticity, a possibility consistent with the improved rotarod performance.

Interestingly, our data indicate that this enhancement in behavioral acquisition may come at the expense of flexible, online behavioral adjustments as illustrated by the sustained engagement in lever pressing during the progressive ratio task. This Smo-associated trade-off suggests the existence of mechanisms that carefully calibrate the balance of Smo and DA signaling on CIN.

### Smo activity on CIN is critical for CIN plasticity

The triphasic activity pattern of CIN, consisting of an initial burst, a pause, and a rebound, dynamically adapts across the course of learning.^10^ In particular, the cholinergic pause initially lengthens with conditioning and then diminishes with overtraining once contingencies are well established. ^10,67,69^ D2R manipulations on CIN, which bidirectionally alter the magnitude and duration of CIN pauses in a manner that resembles these natural changes, have been linked to learning performance.^23,24^ While this relationship suggests that the pause shortening observed across training could arise from diminishing DA release as conditioning cues become less novel, similar pause shortening also occurs as motor actions become more efficient and temporally precise without explicit DA-inducing rewards.^70^ This indicates that reduced D2R engagement on CIN cannot fully account for the progressive shortening of the pause with overextended training. Further, evidence indicates that cholinergic neurons themselves can undergo Hebbian plasticity.^64,71,72^ Consistent with this idea, reduction in behavioral flexibility in response to repeated alcohol exposure is associated with decreased AMPA receptor currents in CIN without accompanying presynaptic changes in release.^9,73^

Here, we observe a robust shortening of pause magnitude and duration across daily sessions of evoked DA burst release, even though the magnitude of the evoked DA response itself remained stable across stimulation days. Notably, this progressive shortening required Smo expression on CIN and was also acutely sensitive to the Smo agonist SAG. Consistent with the dynamics of these changes, Smo is well known to impact neurite maturation and neural growth cone guidance in a progressive manner over hours to days through gene transcription dependent and independent mechanisms.^38,74^ Further, SAG activation of Smo can rapidly reduce cAMP levels, recruit β-arrestin, activate GTPases, and induce intracellular calcium fluctuations through Gαi signaling pathways.^75–78^ Gαi-coupling allows Smo to modulate fast secondary messengers that strongly influence neuronal excitability, suggesting parallel arms of Smo signaling that could support both the acute and the progressive components of ACh pause modulation that we observed.

### Smo dependent signaling within striatal circuitry

Our findings raise questions about how Smo activity in CIN is evoked and constrained in the undisturbed striatum. In this context it is important to point to several lines of evidence suggesting the existence of a cell type specific peptidergic signaling network in the striatum that is based on Smo.

Shh, the ligand that binds to Ptch1 which causes the activation of Smo, is produced by both DAN and PTN.^26,35–37^ Thus, Smo, whose downstream effects are strictly Shh dose dependent in developmental contexts ^38^, could detect coincident Shh release from DAN and PTN providing a possible learning rule that might in part govern the extent of CIN plasticity. Further, CIN elaborate a primary cilium which forms a sub-compartment of the plasma membrane that is critical for transducing Smo-dependent signals to the nucleus and regulating gene expression.^27,30,38^ Notably, primary cilia receive input from neuronal projections, including DAN afferents.^79^ Smo has also been observed at perisynaptic membranes ^41,42^, where it likely acts through Gαi-protein signaling cascades. This subcellular compartmentalization might segregate the long term, and acute, branches of Smo signaling within CIN raising the possibility that Shh release from DAN and PTN influences CIN plasticity with distinct temporal and spatial specificity, depending on the relative engagement of ciliary versus synaptic Smo signaling within the same neurons.

Conversely to the expression of Shh by DAN and PTN, Ptch1is expressed in the striatum by fast-spiking interneurons (FSI) as well as by CIN.^26^ Because FSI provide sparse but powerful GABAergic inhibition of CIN ^80–83^, Shh signaling may influence CIN activity through both direct modulation and indirect modulation via FSI. In this scenario, coincident Shh release onto CIN and FSI could initially attenuate CIN inhibition but subsequently counteract this effect by enhancing FSI-mediated inhibition. Without excluding potential contributions from MSNs or midbrain GABAergic projections ^66^, this local Smo-centered microcircuit could provide a parsimonious explanation for our observation that SAG administration in animals lacking Smo on CIN produced a deepened pause following DAN stimulation: Here, because our experimental design preserved Smo expression on all other striatal cell types, acute SAG treatment selectively revealed the net indirect effects of Smo signaling onto CIN. It is notable that this indirect effect not only opposed the direct action of Smo activation on CIN, but was also stable across repeated stimulations, strengthening our conclusion that changes in pause duration are governed by cell-autonomous actions of Smo on CIN. Furthermore, this activator–delayed inhibition motif resembles classical components of Turing reaction-diffusion systems that can model Shh/Smo signaling in developmental contexts ^84–87^. While highly speculative, it is intriguing to consider that Smo-dependent modulation of FSI-CIN microcircuits could influence the propagation and termination of ACh waves recently identified in the DLS, which themselves display properties consistent with Turing reaction-diffusion mechanisms.^88^

### Implications for Parkinson’s Disease

Recent evidence shows that most individuals with sporadic Parkinson’s Disease exhibit marked deficits in primary cilium maintenance among CIN together with the hallmark severe degeneration of DAN. Rodent models additionally reveal that deficient primary cilia formation on CIN result in reduced expression of dopaminergic trophic factor GDNF and diminished Shh/Smo signaling in CIN.^27–29,89–91^

Together with these clinical and preclinical observations, our results identifying Smo as a critical modulator of DA–ACh coordination during learning begin to suggest that early disruptions in Shh/Smo signaling might impair striatal plasticity before overt neurodegeneration emerges. In this view, physiological fluctuations in Smo activity that normally support adaptive learning may, if dysregulated, contribute to pathological circuit remodeling and disruption of trophic factor signaling, highlighting a potential mechanism linking well established early synaptic dysfunction to later neuronal degeneration.

### Limitations of the study

It is important to note that this neuronal population-based study does not take into consideration functional heterogeneity of CIN. There are at least two waves of striatal cholinergic neuron differentiation during development which produce molecular, and likely physiological, distinct populations of CIN that populate striatal compartments at different ratios.^92–96^ Here we focused on the DLS without distinguishing CIN subtypes and compartmentalization into striosomes and matrix therein. Recent evidence suggests that these DLS sub-compartments might be populated by CIN of distinct physiology.^97^ Further, previous observations revealed that about 50 % of CIN are sensitive to an interruption of Smo signaling in mouse and humans in the DLS.^26,27^ These considerations suggest that the effect sizes of ACh based readouts of Smo manipulations presented here represent likely an underestimation. Further, additional impacts of Smo signaling on CIN might have been masked or diluted by the activity of non Smo dependent CIN.

A further limitation of the current study is the absence of simultaneous cholinergic recordings during behavior. However, the use of the same genetic models in which dorsolateral CIN pause dynamics were quantified, combined with a focus on DLS-dependent behaviors, supports a role for CIN-specific Smo signaling in shaping behavioral output.

## Resource availability

### Lead contact

Requests for further information and resources should be directed to the lead contact, Andreas Kottmann.

### Materials availability

No new materials were generated in this study.

### Data and code availability

The dataset generated supporting this study is available from the lead contact upon request. This paper does not report any original code.

Any additional information required to reanalyze the data reported in this paper is available from the lead contact upon request.

## Acknowledgments

This work was supported by NIH T32GM136499 (PIs: Ruth Stark and Mark Steinberg) through S.U.’s G-RISE fellowship, NIH NIA AG065682 (PI: AHK) and NIH U54MD017979 (PI: Maria Lima), We are grateful to Aria Walls for insightful discussions and technical assistance with fiber photometry data collection and animal care. We thank Sonia Bernal for her essential contributions to laboratory maintenance and organization. We also thank Anis Choudhury, Nicholas Cordero, and Mohammed Kadir for their help with behavioral data collection. Technical support for the fiber photometry systems and data analysis pipeline was generously provided by Christoph Kellendonk, Nicolas Tritsch, and Dustin Zuelke. We further thank Christoph Kellendonk and Andrew Delamater for their helpful feedback on early versions of the manuscript.

## Author contributions

Conceptualization, S. U. and A.H.K.; formal analysis, S.U.; funding acquisition, A.H.K.; investigation, S.U.; writing – original draft, S.U.; methodology, S.U. and A.H.K. writing – review and editing, A.H.K.; visualization, S.U.; software, S.U.; resources, A.H.K.; supervision, A.H.K.

## Supplemental information

Document S1. Figures S1–S7.

## Supplementary Information

**Supplementary Figure 1.**
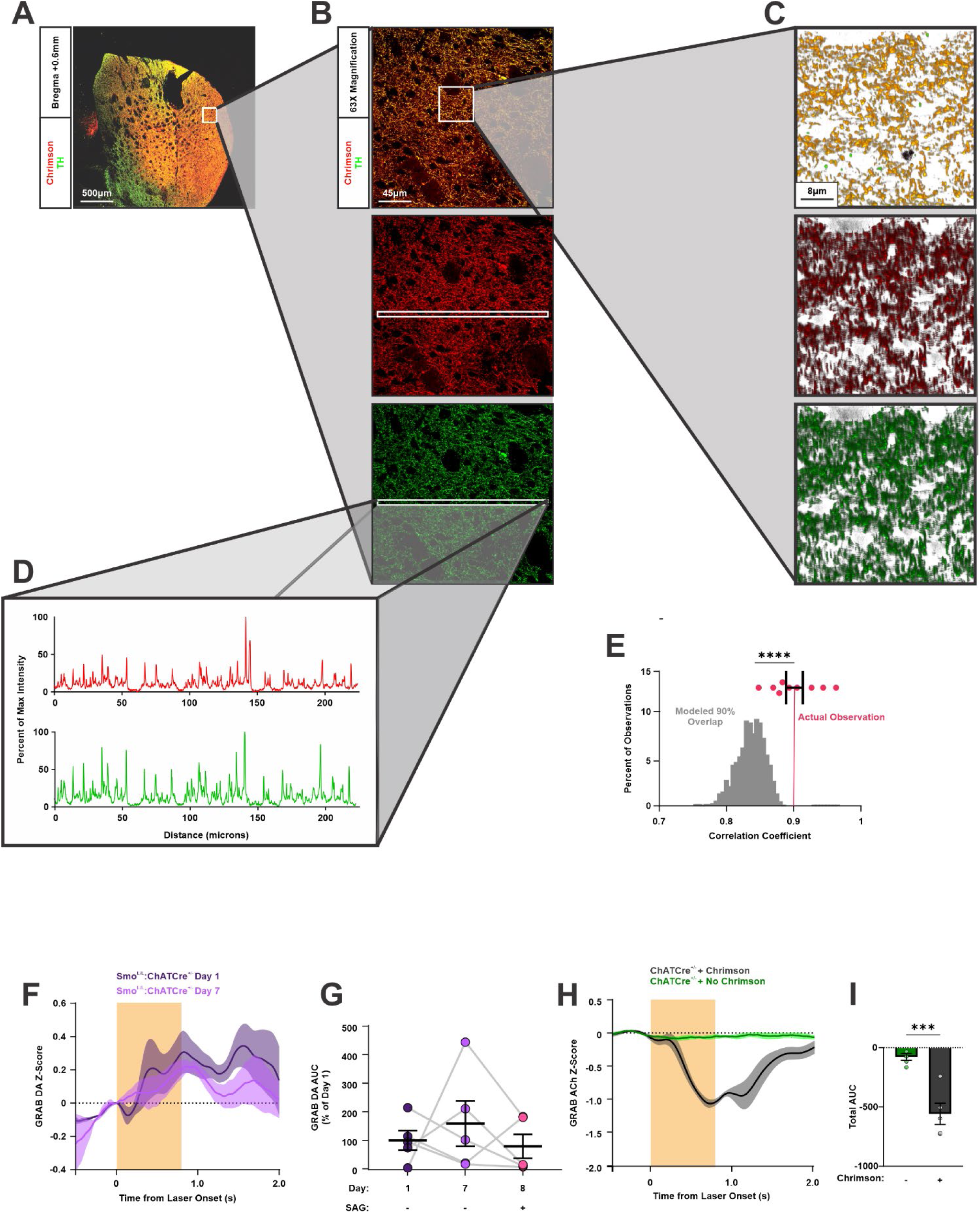
Midbrain Chrimson expression induces Striatal DA release and ACh pauses. **(A)** Representative image showing overlap of Tyrosine Hydroxylase (TH) positive fibers and Chrimson-tdTomato expression across the striatum. Scale bar = 500µm. **(B)** Representative image showing high overlap of TH positive fibers and Chrimson-tdTomato expression at high magnification in the striatum. Isolated TH and Chrimson-tdTomato channels included below. Scale bar = 45µm. **(C)** Z stack projection of subsection in Panel (B) with a depth of 2 µm showing three-dimensional overlap of TH and Chrimson-tdTomato. Isolated TH and Chrimson-tdTomato channels included below. Scale bar = 8µm. **(D)** Pixel intensity for TH and Chrimson-tdTomato fluorescent signals across the one-dimensional arrays highlighted in Panel (B). Pixel intensity was reported as the percentage of maximum intensity observed. **(E)** Comparison between the observed correlation of TH and Chrimson-tdTomato fluorescence signals and a modeled null distribution representing partial (90%) signal overlap. TH and Chrimson-tdTomato overlap was quantified as described in Panel (D), and Pearson correlation coefficients were computed for each striatal section (n = 9). The null distribution was generated by segmenting each Chrimson-tdTomato signal into contiguous 2.5-micron segments, randomly shuffling 10% of segments while keeping the remaining 90% fixed, and recomputing correlations with the corresponding TH signals across 1,000 iterations. The resulting distribution represents correlations expected under a model with 90% overlap between the two signals. The observed mean correlation (r = 0.902) exceeded all values from the simulated null distribution (****p < 0.0001, right-tailed). **(F)** Dopamine (DA) signals aligned to laser onset on days 1 and 7 in Smo^L/L^:ChATCre^+/-^ mice (n = 5). **(G)** Quantification of DA release area under the curve (AUC) from Panel (F) reported as percent of day 1 (n = 5 per day; repeated-measures one-way ANOVA: F(1.443, 5.773) = 0.75, p > 0.05).” **(H)** ACh signal aligned to laser onset from ChATCre^+/-^ mice expressing ACh GRAB sensor with or without midbrain Chrimson expression (n = 5 per Chrimson condition). **(I)** Comparison of GRAB ACh AUC from ChATCre^+/-^ mice in Panel (H) (n = 5 per Chrimson condition; unpaired two-tailed Student’s t test, ***p < 0.001).

**Supplementary Figure 2.**
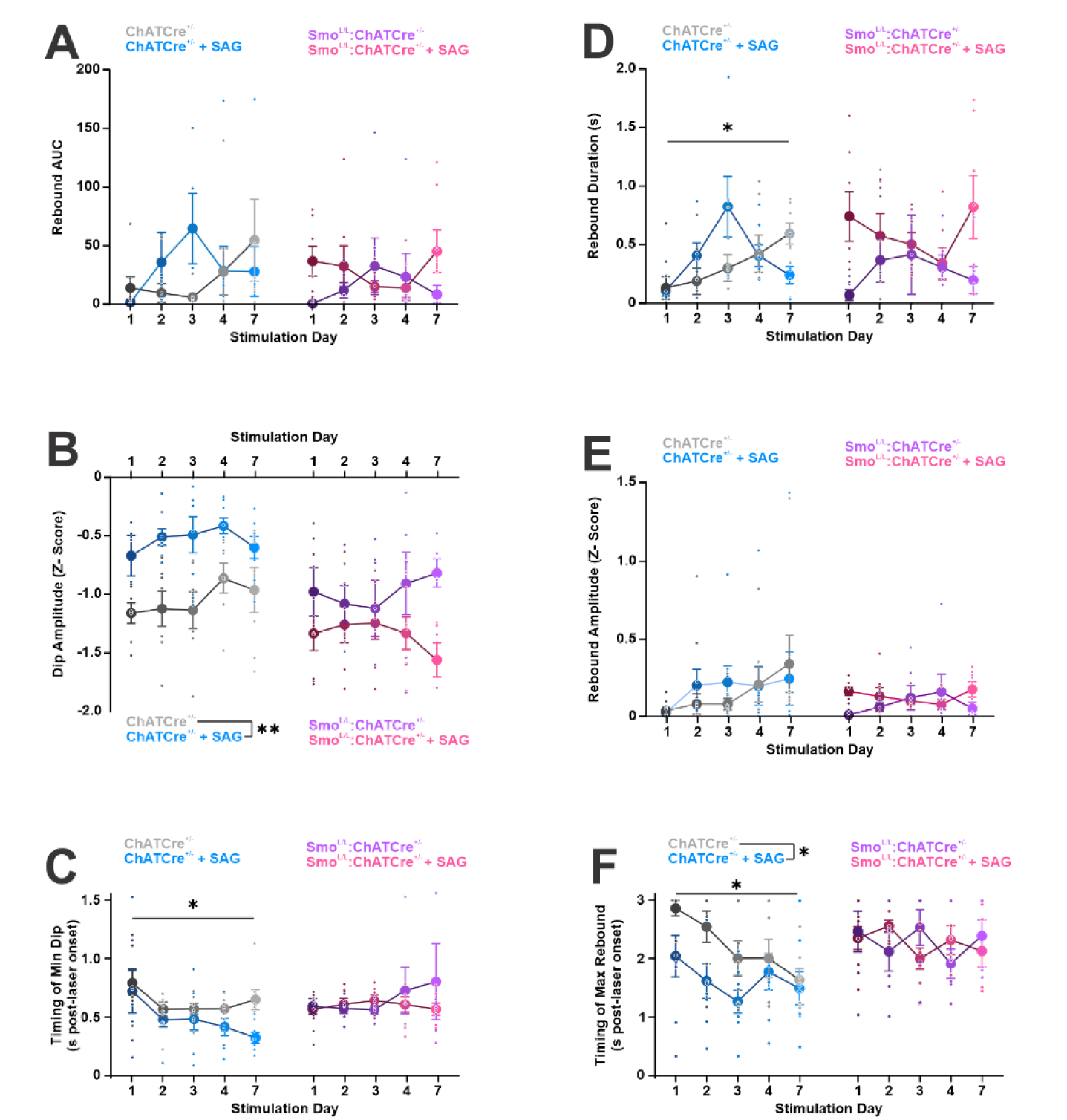
Progression of ACh dip and rebound with repeated DAN stimulation. **(A)** ACh rebound area under the curve (AUC) across stimulation days with or without SAG treatment in ChATCre^+/-^ (n = 7–8 per day; two-way repeated-measures ANOVA: Day effect, F(2.157, 28.04) = 1.49, p > 0.05; Pharmacology effect, F(1, 13) = 0.17, p > 0.05; Day × Pharmacology interaction, F(4, 52) = 2.57, p < 0.05) and Smo^L/L^:ChATCre^+/-^ mice (n = 6–7 per day; two-way repeated-measures ANOVA: Day effect, F(3.295, 36.24) = 0.16, p > 0.05; Pharmacology effect, F(1, 11) = 1.22, p > 0.05; Day × Pharmacology interaction, F(4, 44) = 2.12, p > 0.05). **(B)** ACh dip amplitude across stimulation days with or without SAG treatment in ChATCre^+/-^ (n = 7–8 per day; two-way repeated-measures ANOVA: Day effect, F(2.449, 31.84) = 1.9, p > 0.05; Pharmacology effect, F(1, 13) = 14.12, p < 0.01; Day × Pharmacology interaction, F(4, 52) = 0.65, p > 0.05) and Smo^L/L^:ChATCre^+/-^ mice (n = 6–7 per day; two-way repeated-measures ANOVA: Day effect, F(2.491, 27.41) = 0.16, p > 0.05; Pharmacology effect, F(1, 11) = 2.93, p > 0.05; Day × Pharmacology interaction, F(4, 44) = 3.17, p < 0.05). **(C)** Timing of ACh dip minimum value across stimulation days with or without SAG treatment in ChATCre^+/-^ (n = 7–8 per day; two-way repeated-measures ANOVA: Day effect, F(2.040, 26.51) = 4.00, p < 0.05; Pharmacology effect, F(1, 13) = 3.13, p > 0.05; Day × Pharmacology interaction, F(4, 52) = 0.87, p > 0.05) and Smo^L/L^:ChATCre^+/-^ mice (n = 6–7 per day; two-way repeated-measures ANOVA: Day effect, F(1.800, 19.81) = 0.36, p > 0.05; Pharmacology effect, F(1, 11) = 0.35, p > 0.05; Day × Pharmacology interaction, F(4, 44) = 0.61, p > 0.05). **(D)** ACh rebound duration across stimulation days with or without SAG treatment in ChATCre^+/-^ (n = 7–8 per day; two-way repeated-measures ANOVA: Day effect, F(2.134, 27.74) = 3.41, p < 0.05; Pharmacology effect, F(1, 13) = 0.59, p > 0.05; Day × Pharmacology interaction, F(4, 52) = 3.32, p < 0.05) and Smo^L/L^:ChATCre^+/-^ mice (n = 6–7 per day; two-way repeated-measures ANOVA: Day effect, F(2.729, 30.02) = 0.32, p > 0.05; Pharmacology effect, F(1, 11) = 4.80, p > 0.05; Day × Pharmacology interaction, F(4, 44) = 1.43, p > 0.05). **(E)** ACh rebound amplitude across stimulation days with or without SAG treatment in ChATCre^+/-^ (n = 7–8 per day; two-way repeated-measures ANOVA: Day effect, F(1.383, 17.98) = 3.29, p > 0.05; Pharmacology effect, F(1, 13) = 0.05, p > 0.05; Day × Pharmacology interaction, F(4, 52) = 0.90, p > 0.05) and Smo^L/L^:ChATCre^+/-^ mice (n = 6–7 per day; two-way repeated-measures ANOVA: Day effect, F(2.140, 23.54) = 0.16, p > 0.05; Pharmacology effect, F(1, 11) = 1.10, p > 0.05; Day × Pharmacology interaction, F(4, 44) = 2.21, p > 0.05). **(F)** Timing of ACh rebound maximum value across stimulation days with or without SAG treatment in ChATCre^+/-^ (n = 7–8 per day; two-way repeated-measures ANOVA: Day effect, F(3.327, 43.26) = 4.51, p < 0.01; Pharmacology effect, F(1, 13) = 5.22, p < 0.05; Day × Pharmacology interaction, F(4, 52) = 1.13, p > 0.05) and Smo^L/L^:ChATCre^+/-^ mice (n = 6–7 per day; two-way repeated-measures ANOVA: Day effect, F(2.918, 32.10) = 0.38, p > 0.05; Pharmacology effect, F(1, 11) = 0.01, p > 0.05; Day × Pharmacology interaction, F(4, 44) = 1.45, p > 0.05).

**Supplementary Figure 3.**
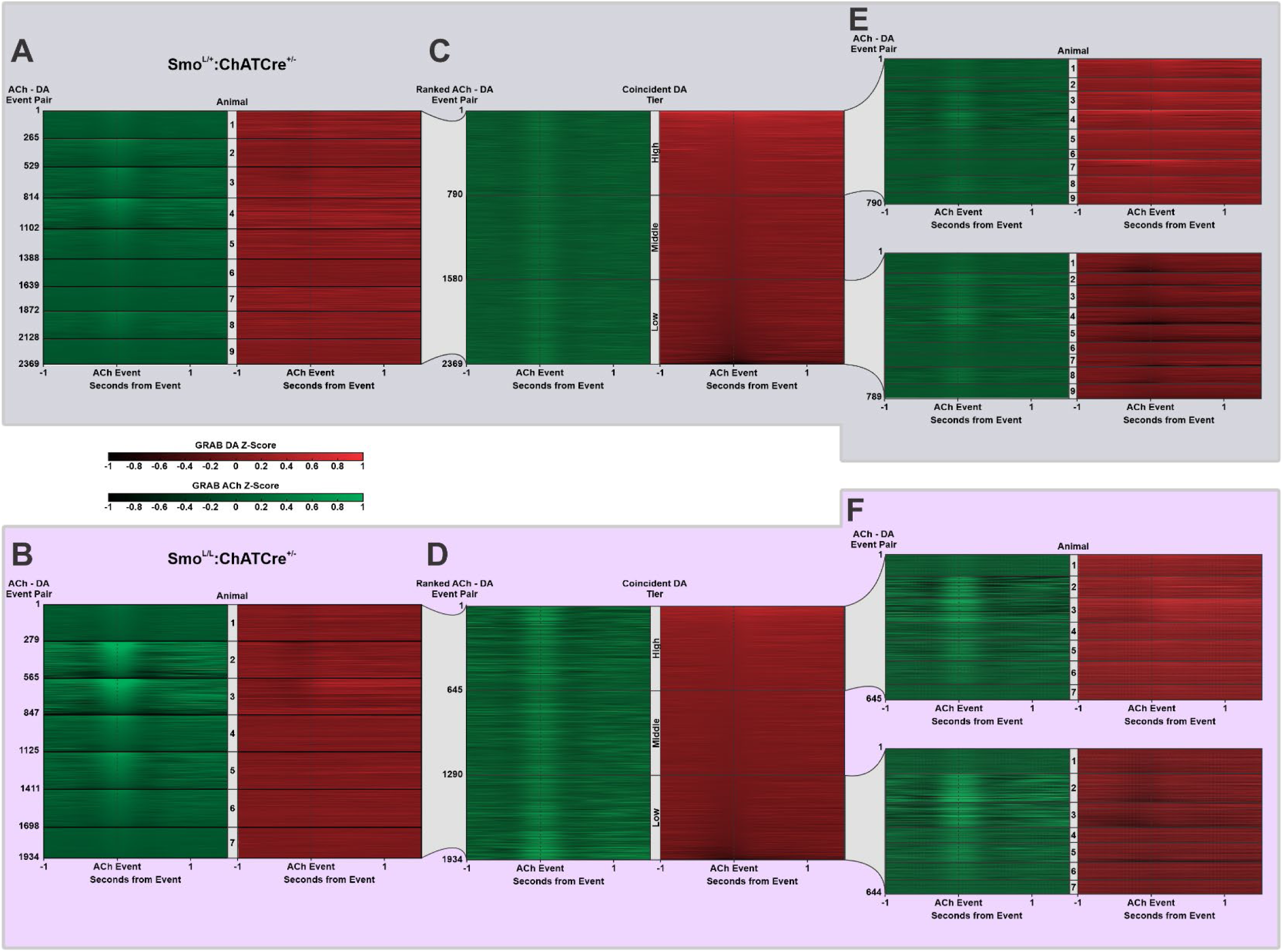
Stratification of ACh Events by levels of coincident DA. **(A)** Acetylcholine (ACh) Events and coincident dopamine (DA) values from control mice, grouped by animal and ranked by ACh Event amplitude. **(B)** Same as Panel (A) for Smo^L/L^:ChATCre^+/-^ mice. **(C)** All ACh Events from control mice, grouped and ranked by the amplitude of coincident DA. Events were stratified into high, middle, and low DA tiers. **(D)** Same as Panel (C), for Smo^L/L^:ChATCre^+/-^ mice. **(E)** ACh Event–DA pairs from high and low DA tiers regrouped by animal to derive new animal-level averages. Within-animal traces are ranked by coincident DA. **(F)** Same as Panel (E) for Smo^L/L^:ChATCre^+/-^ mice.

**Supplementary Figure 4.**
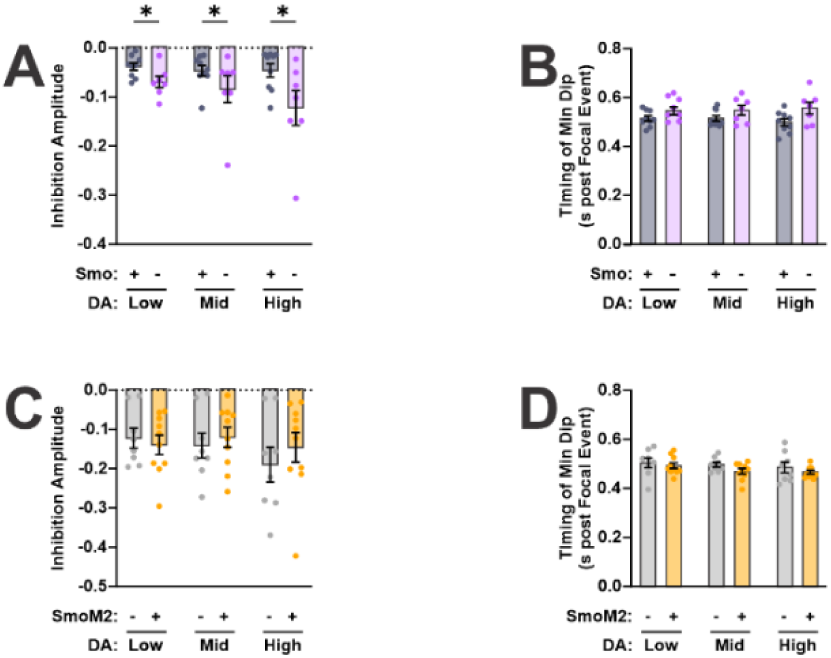
Effects of Smo manipulations on ACh Post-Event Inhibition Amplitude and Timing. **(A)** Amplitude of acetylcholine (ACh) Post-Event Inhibition compared between controls and Smo^L/L^:ChATCre^+/-^ mutants across levels of coincident DA (n = 7–9 per condition; two-way repeated measures ANOVA: Genotype effect, F(1, 14) = 4.79, p < 0.05; Coincident DA effect, F(1.583, 22.16) = 2.95, p > 0.05; Genotype × Coincident DA interaction, F(2, 28) = 1.69, p > 0.05). No post hoc tests were performed due to the absence of a significant interaction effect; graph annotations denote significant main effects only. **(B)** Timing of minimum Post-Event Inhibition compared between controls and Smo^L/L^:ChATCre^+/-^mutants across levels of coincident DA (n = 7–9 per condition; two-way repeated measures ANOVA: Genotype effect, F(1, 14) = 4.30, p > 0.05; Coincident DA effect, F(1.817, 25.43) = 0.08, p > 0.05; Genotype × Coincident DA interaction, F(2, 28) = 1.13, p > 0.05). **(C)** Amplitude of ACh Post-Event Inhibition compared between controls and SmoM2^C/-^:ChATCre^+/-^mutants across levels of coincident DA (n = 8–10 per condition; two-way repeated measures ANOVA: Genotype effect, F(1, 16) = 0.14, p > 0.05; Coincident DA effect, F(1.717, 27.47) = 5.2, p < 0.05; Genotype × Coincident DA interaction, F(2, 32) = 2.76, p > 0.05). **(D)** Timing of minimum Post-Event Inhibition compared between controls and SmoM2^C/-^:ChATCre^+/-^mutants across levels of coincident DA (n = 8–10 per condition; two-way repeated measures ANOVA: Genotype effect, F(1, 16) = 2.85, p > 0.05; Coincident DA effect, F(1.743, 27.89) = 1.34, p > 0.05; Genotype × Coincident DA interaction, F(2, 32) = 0.15, p > 0.05).

**Supplementary Figure 5.**
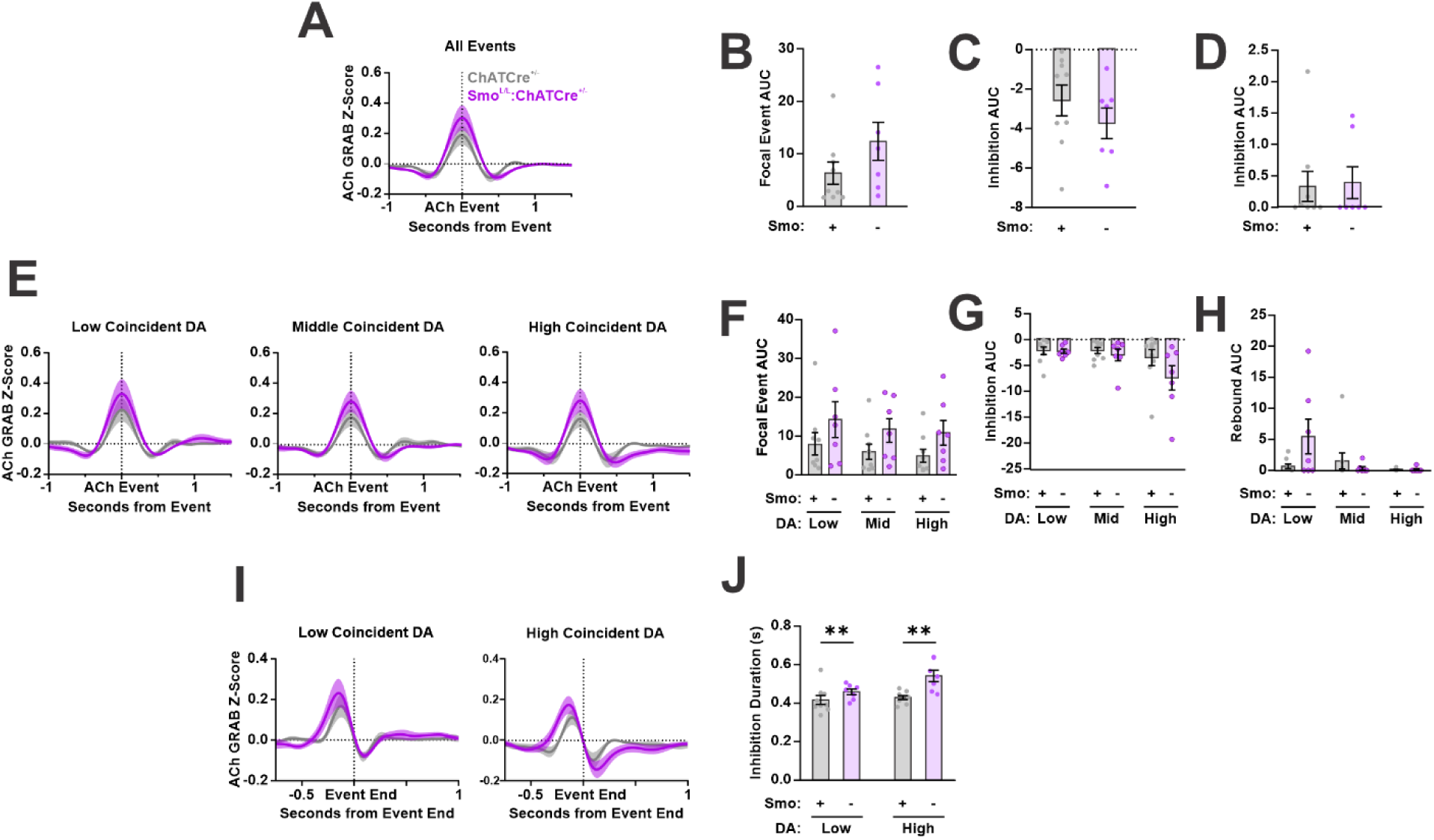
Ablation of Smo from CIN alters DA-associated ACh inhibition relative to alternate homozygous controls. **(A)** Genotype-averaged acetylcholine (ACh) Events for ChATCre^+/-^ control and Smo^L/L^:ChATCre^+/-^ mice (n = 7–9 per genotype). **(B)** Area under the curve (AUC) for ACh Focal Events in Panel (A) (n = 7–9 per genotype; unpaired two-tailed Student’s t test, p > 0.05). **(C)** AUC for ACh Post-Event Inhibition in Panel (A) (n = 7–9 per genotype; unpaired two-tailed Student’s t test, p > 0.05). **(D)** AUC for ACh Rebound in Panel (A) (n = 7–9 per genotype; Shapiro-Wilk test indicated lack of normality, p < 0.001; Mann-Whitney U test, p > 0.05). **(E)** Same ACh Events stratified by relative levels of coincident dopamine (DA) (n = 7–9 per condition). **(F)** AUC for ACh Focal Events in Panel (E) (n = 7–9 per condition; two-way repeated measures ANOVA: Genotype effect, F(1, 14) = 2.31, p > 0.05; Coincident DA effect, F(1.179, 16.51) = 4.43, p < 0.05; Genotype × Coincident DA interaction, F(2, 28) = 0.06, p > 0.05). **(G)** AUC for ACh Post-Event Inhibition in Panel (E) (n = 7–9 per condition; two-way repeated measures ANOVA: Genotype effect, F(1, 14) = 1.50, p > 0.05; Coincident DA effect, F(1.206, 16.88) = 5.76, p < 0.05; Genotype × Coincident DA interaction, F(2, 28) = 1.87, p > 0.05). **(H)** AUC for ACh Rebound in Panel (E) (n = 7–9 per condition; two-way repeated measures ANOVA: Genotype effect, F(1, 14) = 1.49, p > 0.05; Coincident DA effect, F(1.484, 20.78) = 3.42, p > 0.05; Genotype × Coincident DA interaction, F(2, 28) = 3.65, p < 0.05). **(I)** Control and Smo^L/L^:ChATCre^+/-^ ACh Events from Panel (E) realigned to Post-Event Inhibition onset (n = 7–9 per condition). **(J)** Duration of Post-Event Inhibition extracted from traces in Panel (I) (n = 7–9 per condition; two-way repeated-measures ANOVA: Genotype effect, F(1,14) = 12.43, p < 0.01; Coincident DA effect, F(1, 14) = 6.48, p < 0.05; Genotype × Coincident DA interaction, F(1,14) = 3.54, p > 0.05). No post hoc tests were performed due to the absence of a significant interaction effect; graph annotations denote significant main effects only.

**Supplementary Figure 6.**
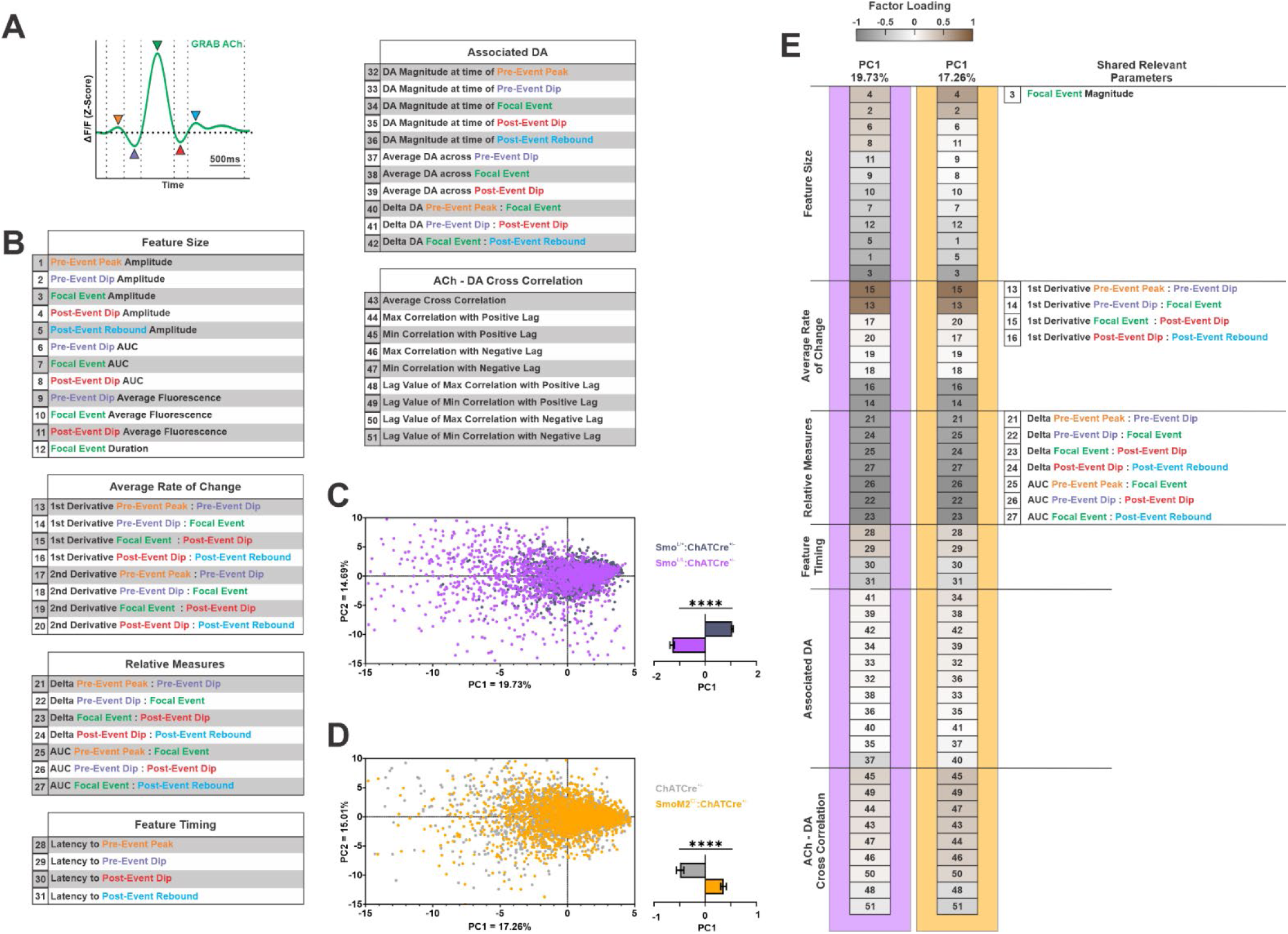
Principal component analysis of ACh Events in Smo loss- and gain-of-function models. **(A)** Schematic of representative acetylcholine (ACh) Event with color-coded arrows denoting features used in analysis and corresponding to Panel (B). All features begin and end at baseline (dashed vertical lines). **(B)** List of parameters used for principal component (PC) analysis, grouped by parameter type. Color-coded labels match features in Panel (A). Colons indicate segments defined between specific features. **(C)** *Left:* PC scores for individual ACh Events from control and Smo^L/L^:ChATCre^+/-^ mice (n = 1926–2357 individual ACh Events per genotype). *Right:* Average PC1 score per genotype (unpaired two-tailed Student’s t test, ****p < 0.0001). **(D)** Same as Panel (C), for control and SmoM2^C/-^:ChATCre^+/-^ mice (n =1827–2445 individual ACh Events per genotype; unpaired two-tailed Student’s t test, ****p <0.0001). **(E)** Parameters correlating with PC1 in datasets comparing control vs. Smo^L/L^:ChATCre^+/-^ (purple) and control vs. SmoM2^C/-^:ChATCre^+/-^ (gold). Parameters are grouped by parameter type and ranked by factor loading. Rightmost column lists features with strong contributions to PC1 in both datasets (|factor loading| > 0.5).

**Supplementary Figure 7.**
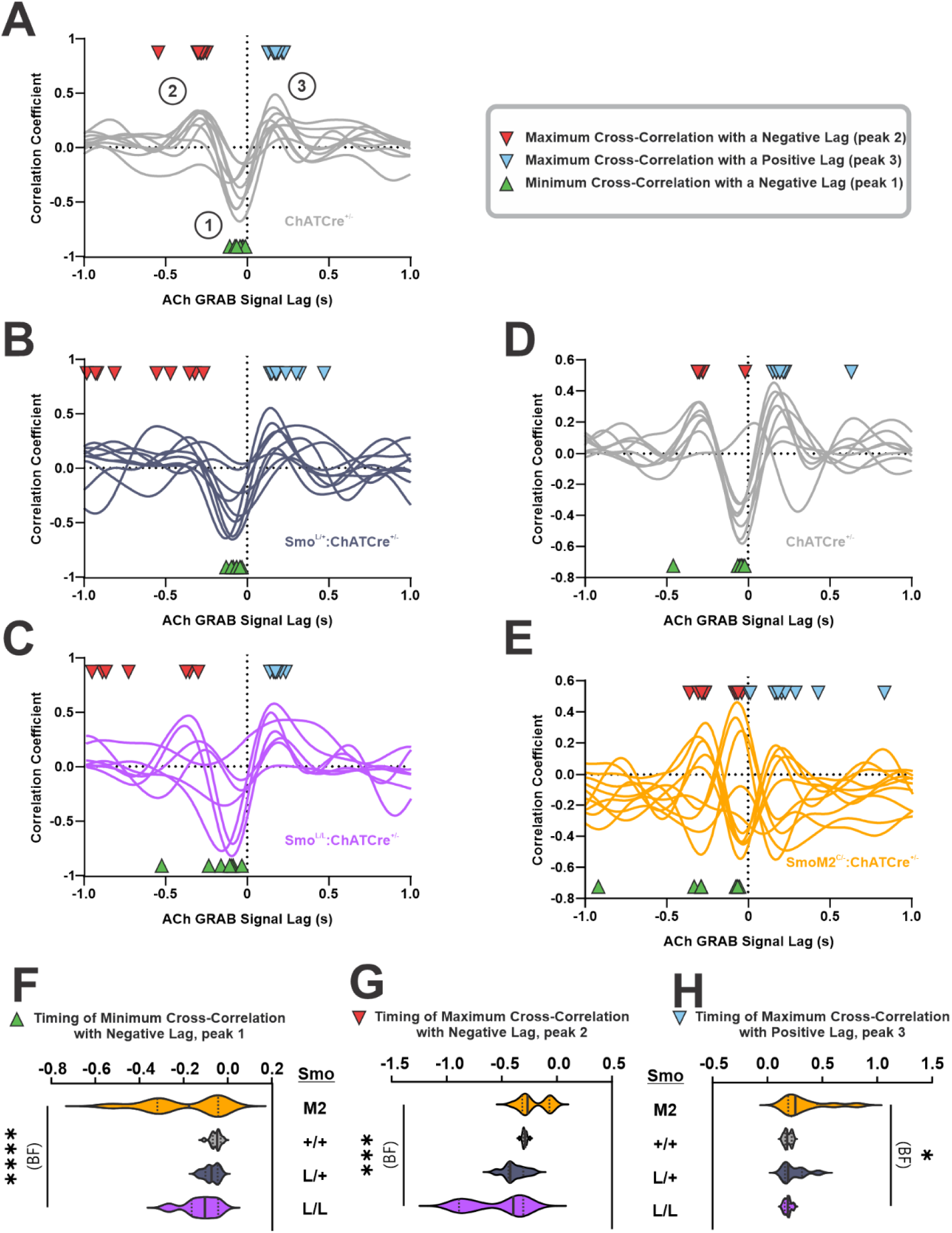
**Variability of cross correlation features across mice** In all Panels: red arrows indicate maximum value of negative lag cross correlation, blue arrows indicate maximum value of positive lag cross correlation, green arrows indicate minimum value of negative lag cross correlation. Average lag values in Figure 5 I-N are derived from cross correlations of individual ACh Event-coincident DA pairs, not these session-average cross correlations. **(A–C)** Session-average cross correlations for each control (A, B) and Smo^L/L^:ChATCre^+/-^ animal **(D–E)** Session-average cross correlations for each control (D) and SmoM2^C/-^:ChATCre^+/-^ animal. **(F)** Violin plots showing the distribution of peak negative correlation lags (negative-lag cases only), with median and interquartile range. Variability across genotypes was compared using the Brown–Forsythe test (n = 7–16 per genotype; Brown-Forsythe test indicated unequal variances, ****p < 0.0001). **(G)** Violin plots showing the distribution of peak positive correlation lags (negative-lag cases only), with median and interquartile range. Variability across genotypes was compared using the Brown–Forsythe test (n = 7–16 per genotype; Brown-Forsythe test indicated unequal variances, ***p < 0.001). **(H)** Violin plots showing the distribution of peak positive correlation lags (positive-lag cases only), with median and interquartile range. Variability across genotypes was compared using the Brown–Forsythe test (n = 7–16 per genotype; Brown-Forsythe test indicated unequal variances, *p < 0.05).

## EXPERIMENTAL MODEL AND STUDY PARTICIPANT DETAILS

All procedures involving animals were conducted in accordance with the National Institutes of Health guidelines and were approved by the Institutional Animal Care and Use Committees of the City University of New York. Experiments were conducted in adult mice between 2 and 8 months of age, weighing between 16 and 34 g. Both male and female mice were used in approximately equal proportions for all experiments. Mice were maintained on a 12 h light/dark cycle with ad libitum access to food and water. Behavioral experiments and recordings were conducted during the light phase with efforts made to balance time-of-day testing across conditions. Animals were group housed for all behavioral experiments. For fiber photometry experiments, animals were group housed prior to surgery and then single housed for the duration of recovery and recording periods.

### Genetic Strategy

Because Smo^L/L^ and SmoM2^C/-^ alleles were backcrossed onto different genetic backgrounds, each genetic model was compared to controls of the same background strain across experiments. Mice with conditional ablation of Smo from ChAT^+^ cells were generated by crossing Smo^L/+^:ChATCre^+/-^ mice with Smo^L/L^ mice, following an initial pairing of ChATCre^+/-^ with Smo^L/L^ on a 129/Sv background. To account for potential off-target effects of introducing the floxed Smo allele and to facilitate the generation of mutant and control littermates, heterozygous Smo^L/+^:ChATCre^+/-^ mice were included as controls in several experiments, as noted in the text. In some cases, ChATCre^+/-^ mice with two intact Smo alleles also served as alternate controls. Mice with conditional ChAT^+^ expression of the constitutively active Smo variant, SmoM2, were generated by crossing SmoM2^C/-^ mice with ChATCre^+/-^mice on a C57BL/6 background. All animals were toe-clipped at 7-14 days of age, and tissue was collected for DNA extraction and subsequent PCR genotyping of the relevant alleles. Forward and reverse primers are listed in the Key Resources Table.

## METHOD DETAILS

### Pharmacology

Smoothened agonist (SAG hydrochloride) was purchased from Biosynth International Inc. and diluted in sterile water to a stock concentration of 10 mg/ml. The solution was aliquoted to avoid repeated freeze-thaw cycles and stored at −20 °C. For experiments, the stock was further diluted in saline so that a bolus of 10 µl per gram of body weight delivered a dose of 20 Mg/Kg.

### Viral Constructs

The AAV5-Syn-ChrimsonR-tdT construct was obtained from Addgene for optogenetic stimulation experiments. For fluorescence-based detection of Acetylcholine and Dopamine, the following viral constructs were obtained from WZ Biosciences: green-fluorescent AAV9-hSyn-Ach3.0 and AAV9-hSynrDA2h (DA4.3h), and red-fluorescent AAV9-hSynrDA1h (rDA2.5h).

### Implant Preparation

Nine-millimeter lengths of 400 µm-diameter optical fiber (FT400UMT; Thorlabs) were scored using a ruby fiber optic scribe (S90R; Thorlabs) and manually pulled to produce clean breaks. Fiber segments were then glued into 6.4mm-long ceramic ferrules (CFLC440; Thorlabs) with two-part epoxy and left to cure overnight. Implants were polished using a series of lapping sheets with decreasing grit sizes until light transduction efficiency reached ≥75%, as tested with a laser.

### Surgical Procedures

Anesthesia for viral injections and fiber implantations was induced using a gas mixture of oxygen (1.5 L/min) and 4% isoflurane (Covetrus). Mice were positioned in a stereotaxic frame (Model 900; David Kopf Instruments) and maintained under anesthesia with 2-3% isoflurane throughout the surgery. A local anesthetic (70 µl bupivacaine) was administered subcutaneously under the scalp prior to incision. At the conclusion of surgery, 0.05 Mg/Kg buprenorphine (Buprenex) was administered subcutaneously, and mice were placed in clean home cages on a heating pad for recovery.

For optogenetic stimulation experiments, 0.5 µl of AAV carrying ChrimsonR-Tdt was injected into the substantia nigra pars compacta at coordinates AP: −3.2 mm, ML: +1.5 mm, DV: −4.1 mm. A second viral injection of 0.5 µl AAV encoding either the green-fluorescent GRAB DA or GRAB ACh sensor was delivered into the dorsolateral striatum at coordinates AP: +0.6 mm, ML: +2.2 mm, DV: −3.0 mm.

For dual-channel GRAB fluorescence recording experiments, a single 1 µl injection containing a 50:50 mixture of red-fluorescent GRAB DA and green-fluorescent GRAB ACh sensors was infused into the dorsolateral striatum at coordinates AP: +0.6 mm, ML: +2.2 mm, DV: −3.0 mm.

In both optogenetic stimulation and dual-channel GRAB fluorescence experiments, fiber implants were placed at coordinates AP: +0.6 mm, ML: +2.2 mm, DV: −3.0 mm, and secured using Metabond® (Parkell, Inc). In dual-channel GRAB fluorescence animals, a small burr hole was also made at lambda, and a custom headbar was affixed with a screw and secured with Metabond® to facilitate head fixation during recording sessions.

On the day following surgery, implant stability was confirmed, topical antibiotic was applied to the surgical site, and a dust cap (CAPL; Thorlabs) was affixed to protect the implant. A minimum of 2.5 weeks was allowed following surgery to ensure full recovery and sufficient viral expression before beginning experiments.

### Optogenetic Experiments

Experiments were conducted in a 43 x 43 cm open-field arena (ENV-515S; Med Associates), isolated from ambient noise and light, and located in a room housing other mouse cages. Animal movements were recorded and analyzed using Noldus EthoVision XT video tracking system. Fluorescence measurements of DA or ACh levels were acquired using a Doric Lenses 5 port Fluorescence Mini Cube (ilFMC5-G3_IE(400-410)_E(460-490)_F(500-540)_O(580-680)_S). The Mini Cube was connected to a Laser Diode Fiber Light Source (LDFLS 465 nm/80 mW; Doric Lenses) via the optogenetics (O) port using an optical patch cord (MFP_200/220/LWMJ-0.22_1m_FC-FCA; Doric Lenses). ChrimsonR excitation, GRAB sensor excitation, isosbestic excitation (405 nm), and signal collection were all performed through a single patch cord (MFP_200/220/LWMJ-0.37_1m_FCM-MF1.25(F)_LAF; Doric Lenses), connected to the animal’s implanted optical fiber via an interconnect (ADAL3; Thorlabs), and to the sample (S) port of the Mini Cube. To reduce torsion and facilitate natural movement, the patch cord was supported by an aluminum wire suspended above the arena.

The Mini Cube, laser diode, signal acquisition, and synchronization with behavioral recordings (via a Noldus USB-IO box) were all controlled and modulated using a National Instruments USB-6343 multifunction I/O device. Signal timing and routing were programmed in WaveSurfer (Janelia), a MATLAB-based data acquisition platform. ChrimsonR stimulations consisted of 780 ms pulse trains delivered every 35 seconds using 590 nm light at 15Hz. Laser power was calibrated to deliver 15 mW/mm^2^ at the tip of the implant. GRAB sensor excitation was performed using 10 ms pulses of 460 nm light at a frequency of 100 Hz, interleaved with isosbestic excitation using 10 ms pulses of 400 nm light at a frequency of 100 Hz. Emitted fluorescence was detected using the Mini Cube’s silicon photodiode sensor (gain set to 10x) and digitized by WaveSurfer at a sampling rate of 1000 Hz.

At the start of each session, the animal’s fiber implant was securely connected to the patch cord, and the animal was then placed in the center of the arena. Prior to the first stimulation session, animals were habituated in the arena for 20-60 minutes with the patch cord attached, but without ChrimsonR stimulation or GRAB sensor recording. On each of the seven stimulation days, animals were first habituated in the arena for 5 minutes before stimulation began. GRAB sensor recording was omitted on days 5 and 6 to preserve signal strength for the final stimulation day. Animals expressing the GRAB DA sensor underwent an eighth stimulation day, during which a challenge dose of SAG was administered to assess its effects on DA release (Figure 1D, E; Supplementary Figure 1F, G). For all SAG-treated animals, intraperitoneal injections were administered 25 minutes prior to placement in the arena, during which animals remained in their home cage. Data were collected from 5 independent cohorts: 2 for GRAB DA experiments and 3 for GRAB ACh experiments.

### Data Processing

Data files (hierarchical data format version 5, HDF5) generated by WaveSurfer were analyzed using custom MATLAB scripts. Interleaved fluorescence and isosbestic signals from each optogenetic stimulation sweep were separated based on their respective excitation wavelength onset parameters. Peripheral samples from each signal onset were trimmed to account for excitation onset and sweep transition artifacts, and the resulting traces were concatenated into two continuous raw time series. True timestamps were retained to allow precise alignment with the onset of ChrimsonR stimulation. Fluorescence and isosbestic signals were then processed with a zero-phase low-pass filter at 20 Hz, followed by downsampling via a moving mean of 200 samples. ΔF/F% was calculated by fitting the isosbestic signal to the fluorescence signal using linear regression, subtracting the fit from the fluorescence signal, and dividing by the fit, then multiplying by 100. The timing of ChrimsonR stimulation (590 nm light) onset was used to extract 4.5-second ROIs from the ΔF/F% signal, anchored 1.5 seconds before the sample most closely aligned with each stimulation onset. Each individual ROI was then z-scored and baseline-normalized by subtracting the average fluorescence across a 0.5-second window preceding ChrimsonR stimulation from the entire ROI. All ROIs were then averaged to produce a session-averaged trace which was smoothed by applying a 100-sample moving mean.

### Feature quantification

Area under the curve (AUC) for GRAB DA signals aligned to ChrimsonR stimulation was calculated for each session-averaged trace. The maximum fluorescence within a 1.5-second window following ChrimsonR stimulation onset was identified as the peak. Flanking inversion points- determined by changes in the sign of the first derivative of the signal and representing the points of maximal difference from the peak- were identified as the start and end of the peak window. The AUC was then calculated using MATLAB’s trapz function.

Overall AUC for GRAB ACh signals aligned to ChrimsonR stimulation were quantified from the full ACh trace starting at the moment of laser, spanning the dip, and ending after the rebound. Dips were identified as the first stretch of negative values following stimulation onset, beginning when the signal first crossed below zero and ending when it returned to zero or the trace ended, whichever came first. Rebounds were defined as the subsequent stretch of positive values, starting immediately after the dip ended and continuing until the signal returned to zero or the end of the trace. If a trace did not exhibit a clear rebound above baseline, rebound magnitude and duration was defined as zero. Rebound timing in these scenarios was also defined as the timing of the maximum value observed after the negative apex of the preceding dip. For both dips and rebounds, duration, amplitude, and timing of the minimum (for dips) or maximum (for rebounds) signal value were extracted.

### Dual-Channel GRAB fluorescence Experiments

Experiments were conducted in a dark, sound attenuating cubicle (ENV-022MD; Med Associates) isolated from ambient noise and light. Fluorescence measurements of red GRAB DA and green GRAB ACh signals were acquired using a Doric Lenses 5 port Fluorescence Mini Cube (iFMC6_IE(400-410)_E1(460-490)_F1(500-540)_E2(555-570)_F2(580-680)_S), paired with a 4 channel LED driver (LEDD, Doric Lenses) and operated via Doric Neuroscience Studio software. Excitation of red and green GRAB sensors, isosbestic (405 nm) excitation, and fluorescence signal collection were all performed through a single patch cord (MFP_400/440/LWMJ-0.37_1m_FCM-MF1.25(F)_LAF; Doric Lenses), which was connected to the implanted optical fiber via a ceramic sleeve (ADAL1; Thorlabs) wrapped in foil. The other end of the patch cord was attached to the sample (S) port of the Mini Cube. The LED driver and signal acquisition were controlled and modulated using a National Instruments USB-6343 multifunction I/O device.

Red GRAB DA sensor excitation using 10 ms pulses of 555 nm light at a frequency of 33 Hz, green GRAB ACh sensor excitation using 10 ms pulses of 460 nm light at a frequency of 33 Hz, and isosbestic excitation using 10 ms pulses of 400 nm light at a frequency of 33 Hz were interleaved. Emitted red and green fluorescence signals were detected using a pair of silicon photodiode Doric Fluorescence Detectors (DFD_FOA_FC), equipped with a spectral bandpass filter: 525/40 nm for green fluorescence and 630/92 nm for red fluorescence. Each detector was operated in DC detection mode with a gain of 10x, and signals were digitized using WaveSurfer at a sampling rate of 1000 Hz.

At the start of each session, the animal was head-fixed atop a running wheel inside the cubicle and the animal’s fiber implant was connected to the patch cord. Prior to the first recording session, mice were habituated on three separate days for 15 minutes each to head fixation and movement on the running wheel, without GRAB sensor recording. On the day of recording, animals were placed in the apparatus and allowed to habituate for 10 minutes before 30 minutes of data collection. Data were collected from 8 independent cohorts: 5 for Smo^L/L^:ChATCre^+/-^ experiments and 3 for SmoM2^C/-^:ChATCre^+/-^experiments.

#### Data Processing

Data files (hierarchical data format version 5, HDF5) generated by WaveSurfer were analyzed using custom MATLAB scripts. Interleaved red fluorescence, green fluorescence, and isosbestic signals were separated based on their respective excitation wavelength onset parameters. Peripheral samples from each signal onset were trimmed to account for excitation onset transients and potential crosstalk between channels. The resulting traces were concatenated into three continuous raw time series and aligned. Fluorescence and isosbestic signals were processed with a zero-phase low-pass filter at 20 Hz, followed by downsampling using a moving mean over 100 samples. ΔF/F% was calculated for both red and green fluorescence signals by fitting the isosbestic signal to each fluorescence signal via linear regression, subtracting the fit from the fluorescence signal, dividing by the fit, and multiplying by 100. The full green and red ΔF/F% traces were then z-scored.

To identify significant DA Events, we applied a two-pass median absolute deviation (MAD) thresholding approach. First, a 30-second moving median and corresponding 30-second moving MAD were computed across the full GRAB DA z-scored trace. Samples exceeding the median plus 2 MADs were clipped, producing a filtered trace. A second 30-second moving median and MAD were then calculated from this filtered trace. DA Events were recognized as discrete segments in the original z-scored trace that exceeded the filtered median plus 3 MADs. The maximal point of each segment was identified as the DA peak and used for event alignment.

For all extracted events, a 6-second ROI was extracted. To prevent overlap between ROIs, subsequent events that overlapped with a preceding ROI were excluded. Corresponding 6-second ROIs from the aligned GRAB ACh z-scored trace were also extracted. Each ROI was baseline-normalized by subtracting the average fluorescence during the first second of the ROI from the entire segment. All ROIs were then averaged to produce session-averaged traces.

To identify significant ACh Events, the same pipeline described above was used, simply with the roles of the GRAB DA and GRAB ACh traces reversed. The maximal point of each segment in this analysis was again used for event alignment and identified as the Focal Event of each ACh Event.

#### DA Event Aligned Feature quantification

Each coincident ACh signal was decomposed into a burst and dip feature identified by detecting sign changes in the first derivative of the trace.

- Burst Event – the most proximal positive inflection point in the ACh trace preceding the peak of the DA Event.
- Dip Event – the most proximal negative inflection point in the ACh trace following the peak of the DA Event.

Feature boundaries were defined as the first baseline crossings (i.e., points where the signal returns to zero) flanking each identifying inflection.

#### ACh Event Aligned Feature quantification

Each ACh Event was similarly decomposed into a standardized set of features organized relative to the Focal Event (see Supplementary Figure 7A). Inflection points flanking the Focal Event were identified by again detecting sign changes in the first derivative of the trace. ACh Event features were defined as follows:

- Focal Event – the peak of the ACh Event, centered on the defining inflection point.
- Pre-Event Dip – the first inflection point preceding the Focal Event such that the absolute difference in magnitude between this dip and the next most distal inflection point is > 10% of the absolute difference in magnitude between this dip and the Focal Event.
- Pre-Event Peak – the inflection point immediately preceding the Pre-Event Dip.
- Post-Event Dip - the first inflection point following the Focal Event such that the absolute difference in magnitude between this dip and the next most distal inflection point is > 10% of the absolute difference in magnitude between this dip and the Focal Event.
- Post-Event Rebound - the inflection point immediately following the Post-Event Dip, or the end of the trace, whichever occurs first.

Feature boundaries were again defined as the first baseline crossings flanking each identifying inflection. For each feature, the following measures were extracted: magnitude, timing relative to Focal Event, duration, delta (i.e., change in magnitude relative to the Focal Event), AUC relative to baseline, AUC relative to the position of flanking features, first derivate, second derivative, and average fluorescence.

#### Trace Stratification and Realignment

Stratification of DA Events or ACh Events based on coincident DA was performed by grouping all traces from a given genotype, then ranking them by the magnitude of the stratifying measure. The ranked traces were then split into thirds (high, middle, low tiers), and traces within each tier were averaged based on the animal they originated from. For realignment of ACh Events, (Figure 2I) individual ACh Events within each tier of DA-stratified events were aligned to the start of the Post-Event Dip (defined as the final baseline point prior to the inflection point identified as the Post-Event Dip). Animal-level averages were then produced for each tier.

### Behavioral Experiments

#### Rotarod

Trials were conducted in a room with low ambient light and noise using a five-lane mouse rotarod (ENV-574M; Med Associates) programmed to accelerate linearly from 4 to 40 RPM over 300 seconds (preset speed setting #9). After reaching top speed, the trial continued for an additional 400 seconds to assess sustained physical performance and motor skill retention. Mice underwent 10 trials per day for 8 consecutive days. Latency to fall was recorded for each trial. A fall was defined as either falling off the rod or clinging and completing a full passive rotation around the rod. In the latter case, the mouse was manually removed and placed in the bottom of the apparatus. Mice that remained on the rod for the full 700 were also manually removed and placed in the bottom. Mice were allowed approximately 3-5 minutes of rest between trials. As described in the text, the learning criterion was defined as the first day on which a mouse completed the acceleration phase (>300 seconds latency to fall) in more than trials. Data were recorded by an experimenter blinded to genotype and collected from 8 independent cohorts: 5 for Smo^L/L^:ChATCre^+/-^ experiments and 3 for SmoM2^C/-^:ChATCre^+/-^experiments.

#### Lever Pressing

Operant training was performed in a Med Associates modular chamber (ENV-008) with a steel grid floor (ENV-307A-GFW; Med Associates), located in a room housing other mouse cages but isolated from ambient noise. A reward magazine (ENV-303M-4.25; Med Associates) equipped with a head entry detector (ENV-303HDA; Med Associates) was positioned at mouse eye-level in the center of one wall, with a retractable lever (ENV-312-3M; Med Associates) located to the left of the magazine. Chocolate-flavored sucrose pellets (F05301; Bio-Serv) were used as rewards and delivered via an automated pellet dispenser (ENV-203; Med Associates). All equipment was controlled via a Noldus USB-IO box programmed using the Noldus EthoVision XT behavior tracking system.

Animals were first food-restricted to 85% of their baseline body weight and maintained at this weight throughout the experiment by allowing ad libitum access to chow for one hour following each daily lever pressing session. Each session began with a 5-minute habituation period, after which the lever extended and remained available for the full 1-hour session. Head entries, lever presses, and reward deliveries were recorded during all sessions.

Training began with a continuous reinforcement (CRF) phase, during which the magazine and lever were baited with pellet crumbs at the start of the session, and every lever press resulted in a reward. To advance from CRF training, animals were required to complete a minimum of three training days and reach a criterion of 50 presses within a single session—the maximum number of rewards available during this phase. Animals that did not reach this threshold within the first three days continued CRF training until the criterion was met before progressing to the next reinforcement schedule.

Following CRF training, animals underwent Variable Interval 15 (VI15) training, in which an average interval of 15 seconds (range: 0-30 s) had to elapse before lever presses elicited subsequent rewards. After three days of VI15, animals underwent a 15-minute devaluation probe. Prior to the session they were given ad libitum access to either chocolate-flavored sucrose pellets (devalued condition) or standard lab chow (baseline condition). Animals were then placed in the chamber, and lever presses were recorded in the absence of reward delivery.

Animals subsequently completed 8 days of Variable Interval 30 (VI30) training, in which an average of 30 seconds (range: 0-60 s) had to elapse before a lever press was rewarded. For Progressive Ratio 2 (PR2) testing, animals were first required to complete 3 days of CRF, 3 days of VI15, and 6 days of VI30 training, as described above. In the PR2 paradigm, the first reward was delivered after a single press, and the number of presses required for each subsequent reward doubled. PR2 sessions lasted 2 hours and were prematurely terminated if 2 consecutive minutes elapsed without any lever presses. Data for all experiments were pooled from 2 independent cohorts.

## QUANTIFICATION AND STATISTICAL ANALYSIS

All statistical analyses were performed using GraphPad Prism version 10.0.0 for Windows (GraphPad Software, Boston, Massachusetts USA; www.graphpad.com). Statistical tests used, group sizes, p values, and other relevant details are reported in the figure legends. The marker “ns” indicates results that were not significant. Data were assessed for normality (Shapiro–Wilk test) and for homogeneity of variance (Brown–Forsythe or F test). Unless otherwise indicated, all datasets satisfied the assumptions for parametric analysis; when assumptions were violated, the corresponding test results are reported, and nonparametric tests were applied.

All figures display group averages ±SEM. Fiber photometry data processing, analysis, and feature quantification were performed using MATLAB version 9.12.0 (R2022a) for Windows (The MathWorks Inc., Natick, Massachusetts USA; www.mathworks.com).

### Key Resources Table

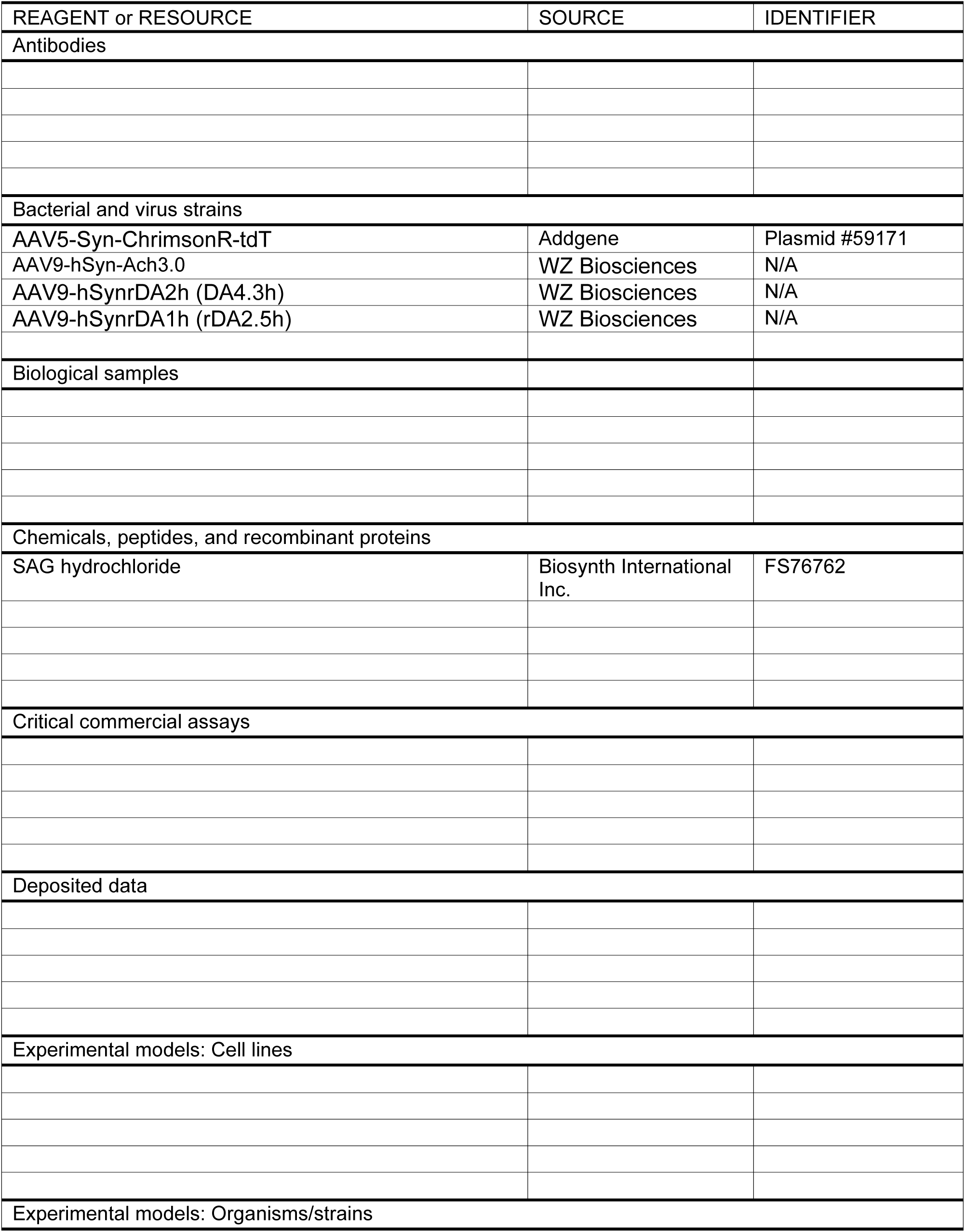
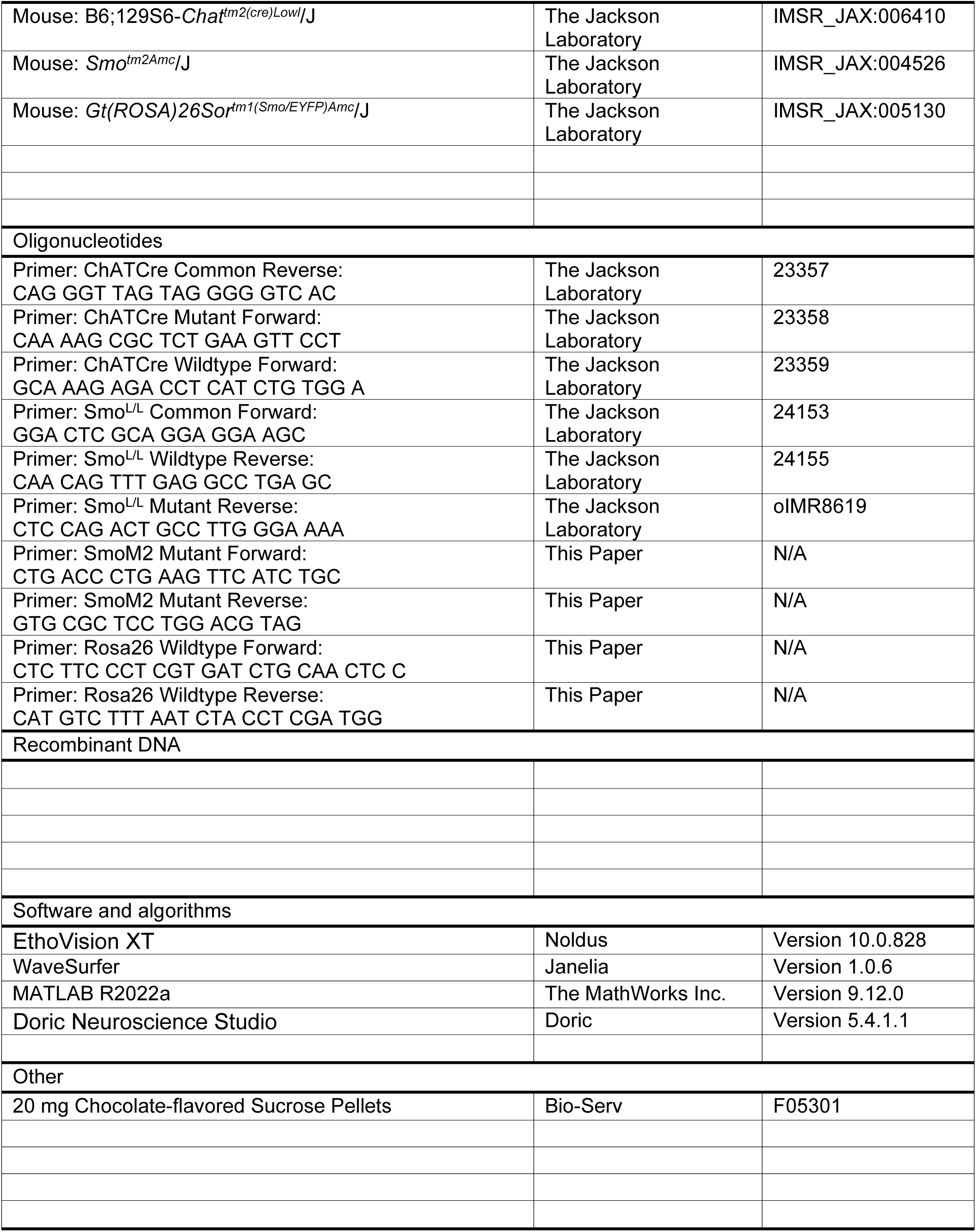

