## supplemental figures 1 - 7 for "The GPCR Smoothened on Cholinergic Interneurons Modulates Dopamine-associated Acetylcholine Dynamics and Affects Learning"

**Supplementary Figure 7. Variability of cross correlation features across mice**
In all Panels: red arrows indicate maximum value of negative lag cross correlation, blue arrows indicate maximum value of positive lag cross correlation, green arrows indicate minimum value of negative lag cross correlation. Average lag values in Figure 5 I-N are derived from cross correlations of individual ACh Event-coincident DA pairs, not these session-average cross correlations.
**(A–C)** Session-average cross correlations for each control (A, B) and $Smo^{L/L}$:$ChATCre^{+/-}$ animal.
**(D–E)** Session-average cross correlations for each control (D) and $SmoM2^{C/-}$:$ChATCre^{+/-}$ animal.
**(F)** Violin plots showing the distribution of peak negative correlation lags (negative-lag cases only), with median and interquartile range. Variability across genotypes was compared using the Brown–Forsythe test (n = 7–16 per genotype; Brown-Forsythe test indicated unequal variances, ****p < 0.0001).
**(G)** Violin plots showing the distribution of peak positive correlation lags (negative-lag cases only), with median and interquartile range. Variability across genotypes was compared using the Brown–Forsythe test (n = 7–16 per genotype; Brown-Forsythe test indicated unequal variances, ***p < 0.001).
**(H)** Violin plots showing the distribution of peak positive correlation lags (positive-lag cases only), with median and interquartile range. Variability across genotypes was compared using the Brown–Forsythe test (n = 7–16 per genotype; Brown-Forsythe test indicated unequal variances, *p < 0.05).

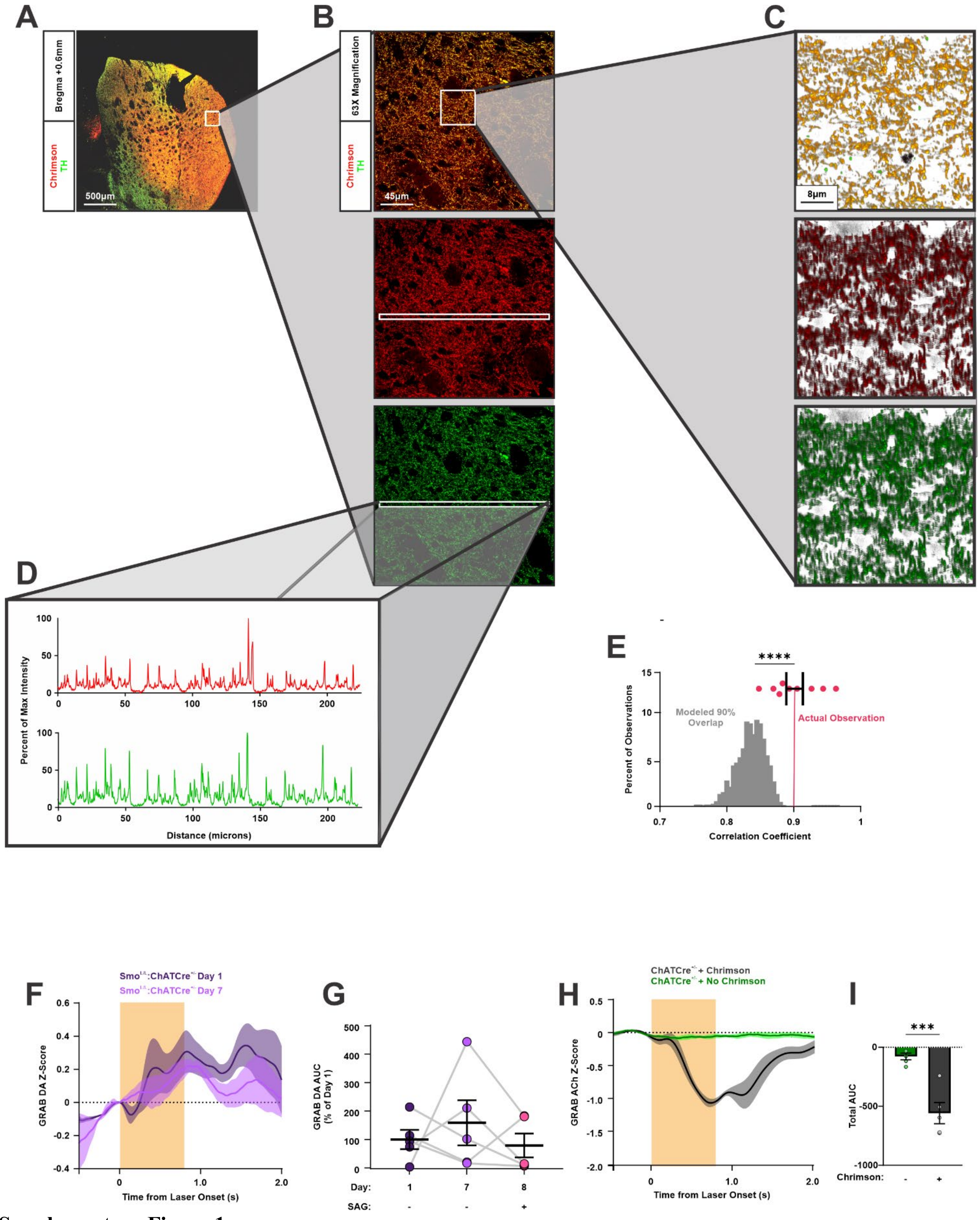


Supplementary Figure 1

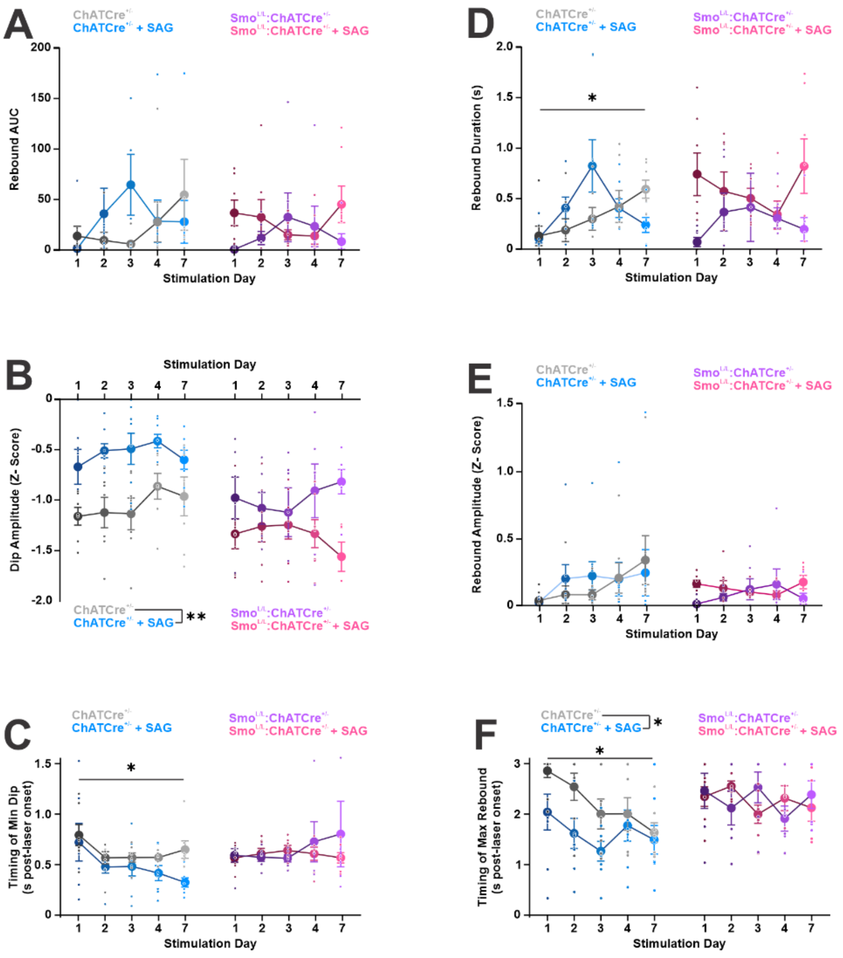


**Supplementary Figure 2**

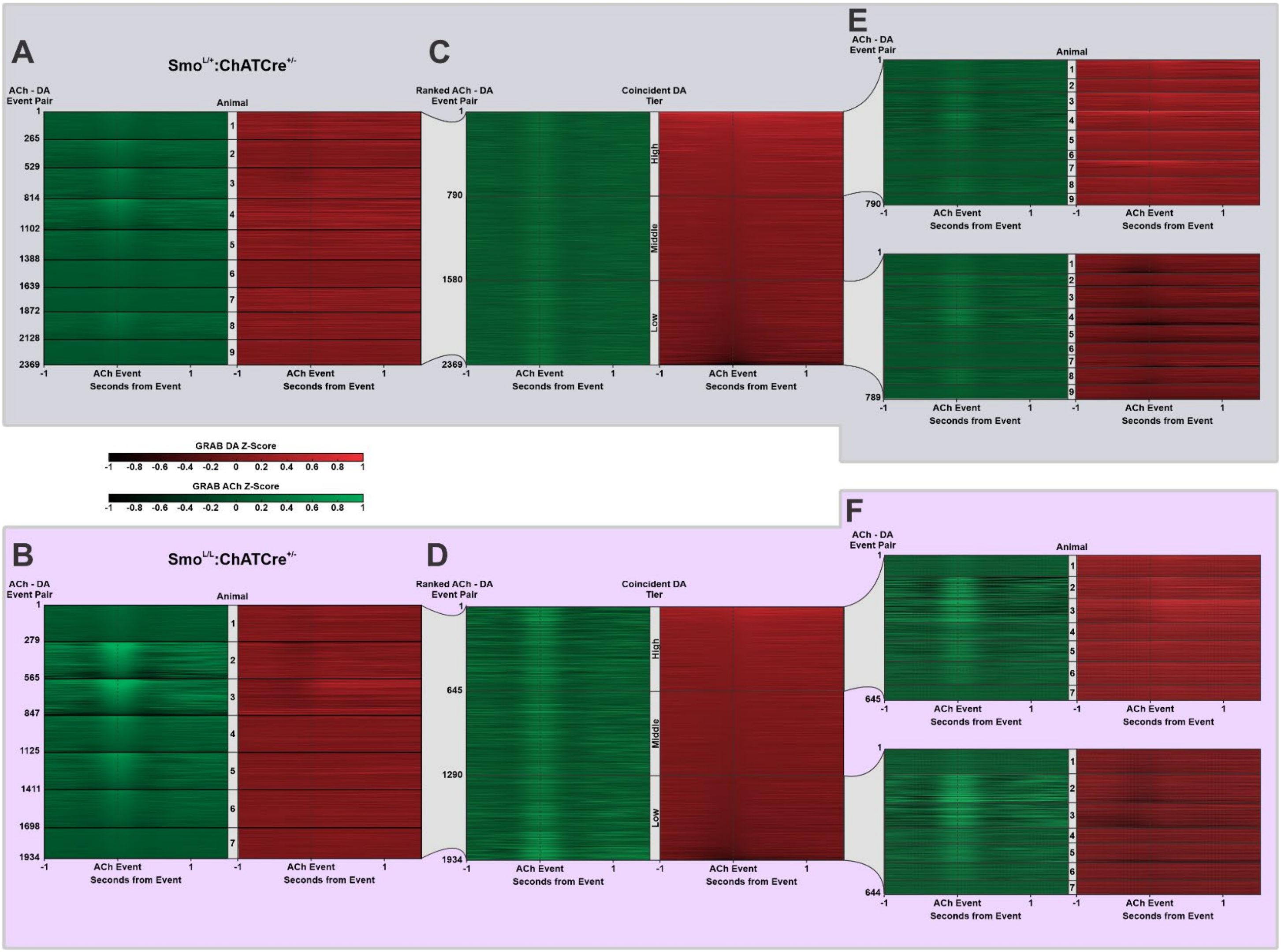


**Supplementary Figure 3**

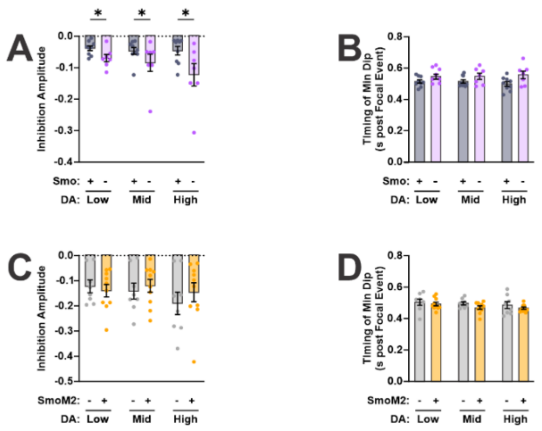


**Supplementary Figure 4**

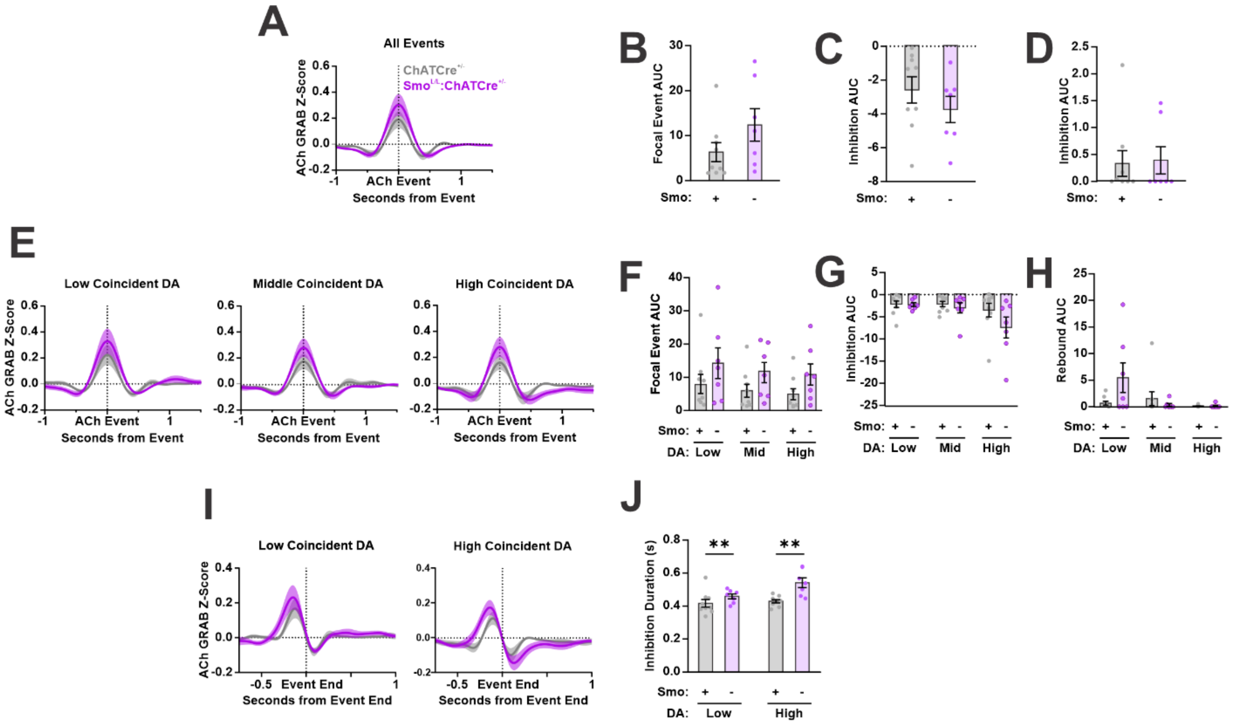


**Supplementary Figure 5**

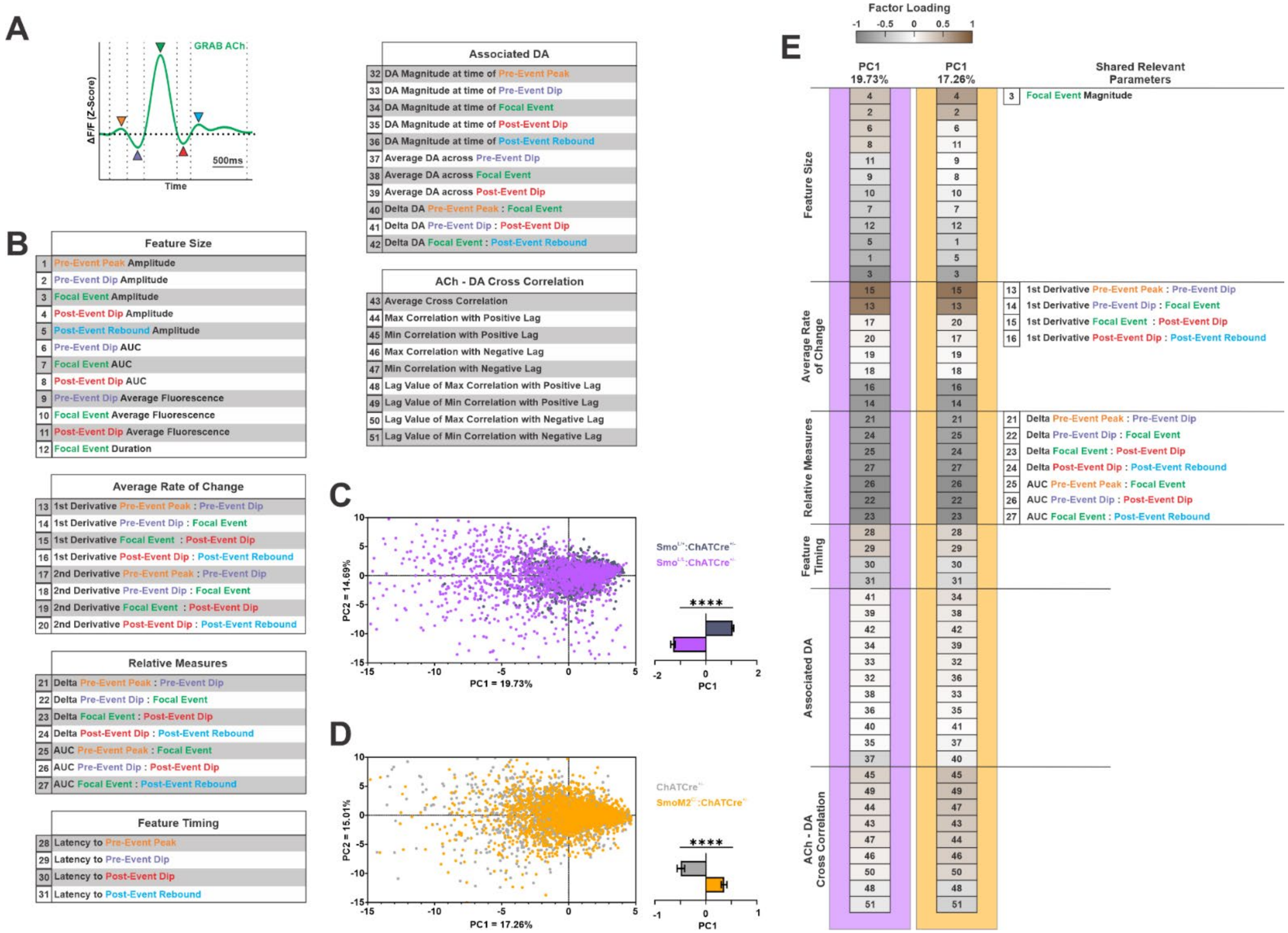


**Supplementary Figure 6**

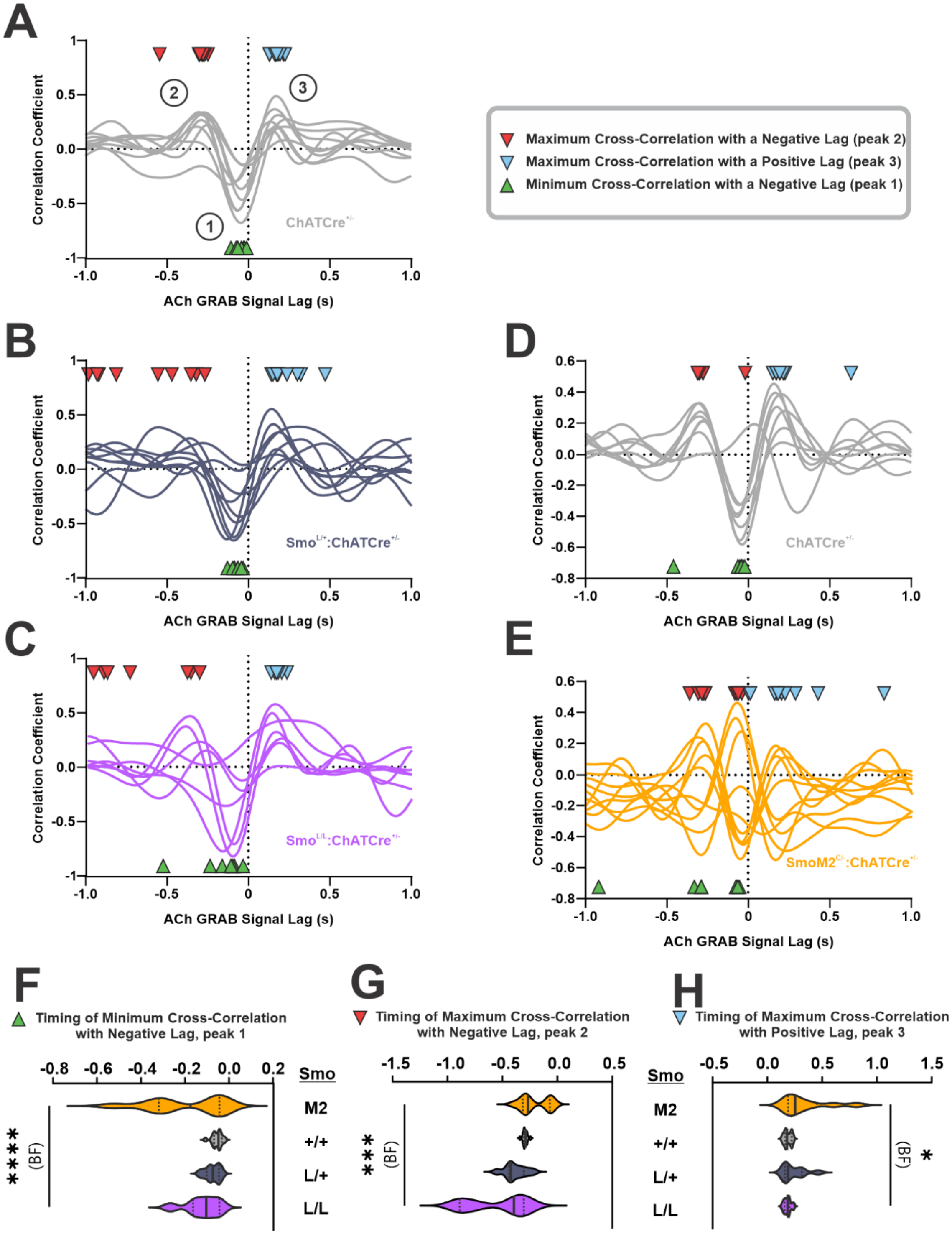


**Supplementary Figure 7**
